# Non-uniform effects of remaining field spread on the estimation of M/EEG activity and connectivity between regions of interest

**DOI:** 10.64898/2025.12.09.688708

**Authors:** Nikolai Kapralov, Alina Studenova, Rubén Eguinoa, Guido Nolte, Stefan Haufe, Arno Villringer, Vadim Nikulin

## Abstract

In M/EEG analyses, it is often convenient to extract time series of activity originating from the regions of interest (ROIs). However, due to the spread of electric and magnetic fields, M/EEG recordings capture activity from all sources within the brain. Commonly used approaches for the extraction of ROI activity only partially alleviate this problem, and field spread remains a challenge even at the level of ROI time series. Because of the remaining field spread (RFS), the extracted time series captures activity not only from the considered ROI but also from other regions (not necessarily neighboring ones). The amount of RFS can strongly affect the validity of interpretations: with more RFS, the extracted time series becomes less representative of the ROI. However, neither the amount nor the pattern of RFS is usually known. In this study, we apply the cross-talk function (CTF) to analyze contributions of all sources across the brain to the extracted ROI time series, thereby quantifying the degree of RFS. With CTF, we show that the effect of RFS on the extraction of ROI activity and on the estimation of inter-regional connectivity is highly non-uniform across the cortex. In particular, ROIs farther away from the recording sensors are more likely to capture activity and connectivity from other areas. Finally, we validate this observation in simulations and complement it by investigating spurious and ghost interactions in real data. Overall, our results illustrate how CTF can be used as a diagnostic tool to quantify the effects of RFS and to evaluate pipelines for the extraction of ROI activity.

## 1. Introduction

The results of neuroimaging analyses are often interpreted and communicated by referring to specific areas of the brain defined according to various parcellations, which are based on anatomical (e.g., Desikan et al. (2006); Destrieux et al. (2010)), functional (e.g., Schaefer et al. (2018)), and cytoarchitectonic (e.g., Brodmann (1909)) criteria, or a combination of those (e.g., Glasser et al. (2016)). In electro- and magnetoencephalography (M/EEG), it is often convenient to define regions of interest (ROIs) and extract time courses of M/EEG activity originating from these ROIs. Such ROI-based representation reduces data dimensionality, making large-scale analyses computationally feasible. Extraction of ROI time series is widely used in basic, cognitive and clinical neuroscience for the analyses of oscillatory power (da Silva Castanheira et al., 2021), evoked responses (Kumral et al., 2022), functional connectivity (Pellegrini et al., 2023; Brkić et al., 2023; Kapralov et al., 2024), biophysical modeling of neural data (Neymotin et al., 2020), and real-time estimation of neural activity for brain stimulation and neurofeedback (Grosse-Wentrup et al., 2009; Zrenner et al., 2018).

Due to the field spread, sensor-space M/EEG recordings contain a mixture of activity from all brain sources (Sarvas, 1987), and inverse modeling methods are commonly used to reconstruct activity from individual brain sources. Still, the reconstruction is never perfect due to the ill-posed nature of the inverse problem (Schoffelen and Gross, 2009; Palva and Palva, 2012). As a result, even after source reconstruction, the extracted ROI time series also captures activity from other brain regions. We refer to this phenomenon as remaining field spread (RFS) throughout the paper.

RFS may crucially affect the validity of our interpretations. Ideally, we aim to attribute the results of all analyses conducted using the extracted ROI time series (e.g., power or connectivity) to the considered ROI. However, with larger RFS, the extracted time series become less representative of the corresponding ROI, and the observed effects might originate from other regions as well. In connectivity analyses, RFS has two well-known consequences – spurious and ghost interactions. Spurious interactions occur when the same sources contribute to the time courses of investigated ROIs. The time courses may thus appear synchronized even if there is no genuine underlying interaction (Nunez et al., 1997; Nolte et al., 2004). In contrast, ghost interactions occur when a genuine interaction between two ROIs can also be observed between other ROIs due to RFS (Palva et al., 2018). Spurious and ghost interactions pose a serious challenge to the interpretation of results, but they can be accounted for by modeling the RFS.

The amount of RFS is generally not known. When RFS is ignored during interpretation, it is implicitly assumed that its effects are either negligible or similar across all brain areas, which may not be true. Intuitively, the amount of RFS should decrease with the distance between the locations where the signals are recorded or reconstructed. Nunez et al. (1997) showed such a decrease in sensor space, and a similar behavior can be expected for ROIs in source-space analysis. However, this idea still doesn’t allow one to determine the amount of RFS more precisely for a particular analysis and interpretation. A better understanding of the RFS could be instrumental in ensuring the correct interpretation of the results.

The properties of linear methods for source reconstruction can be analyzed by computing their resolution matrix (Backus and Gilbert, 1968; de Peralta Menendez et al., 1996). It is common to focus on the rows and columns of the resolution matrix, which are referred to as the cross-talk (CTF) and the point-spread function, respectively (Hauk et al., 2011). CTF shows which ground-truth sources of activity potentially contribute to the reconstructed estimate at a specific location, thereby quantifying the degree of RFS. It was used previously to compare different inverse operators (de Peralta Menendez et al., 1996; Hauk et al., 2011), evaluate the benefit of simultaneous M/EEG recordings for source reconstruction (Molins et al., 2008), and to optimize brain parcellations (Farahibozorg et al., 2018). However, the differences between existing pipelines for extracting ROI activity and the underlying RFS have not yet been analyzed using CTF.

In the present study, we apply CTF to estimate the expected amount and pattern of the RFS for different ROIs and linear extraction pipelines. We first show that every combination of linear methods for extracting ROI activity can be represented by an equivalent spatial filter and associated CTF. We then show how the expected effects of the RFS on the extraction of ROI activity and on the estimation of connectivity between ROIs can be quantified using the CTF. Using simulated data, we further investigate whether CTF can explain the differences in the extraction of activity between brain regions and extraction pipelines. Finally, we show how CTF can explain the spurious interactions observed in resting-state EEG data and ghost interactions observed during steady-state visual stimulation. To facilitate the usage of CTF, we also share the implementation of all key derivations as an open-source Python package ROIextract.

## 2. Theory

This section presents the mathematical derivations of the main ideas presented in the paper. Throughout the section, normal letters (*N_C_*, *r̂*) denote scalars, bold lowercase letters (**x**) – vectors, bold uppercase letters (**L**) – matrices. R denotes the set of real numbers, while E stands for the expected value. The circumflex (”hat”; *r̂*, **ŝ**) is used to denote estimated values. Superscript T denotes transposed matrices. The complete notation is presented in Table S2.

### 2.1. Cross-talk function

The cross-talk function is closely related to the concept of resolution matrix, which shows how well a linear inverse operator can resolve the modeled sources of activity (Backus and Gilbert, 1968; de Peralta Menendez et al., 1996). Let **x**(*t*) ∈ ℝ*^N_C_^*^×1^ and **s**(*t*) ∈ ℝ*^N_C_^*^×1^ be the data recorded from *N_C_* channels and ground truth activity of *N_S_* source dipoles at the time point *t*, respectively. In what follows, we omit the time dependence for conciseness. Assuming that the dipoles are oriented along the normal to the cortical surface, the lead field matrix **L** ∈ ℝ*^N_C_^*^×^*^N_S_^* describes the theoretical relationship between the sensor-space data **x** and source activity **s** in the absence of measurement noise:

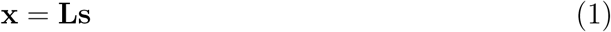

To obtain the reconstructed estimates of source activity **ŝ** ∈ ℝ*^N_S_^* ^×1^ given the sensor-space data **x**, we apply an inverse operator **W** ∈ ℝ*^N_S_^* ^×^*^N_C_^* :

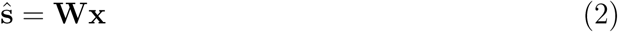

In the equation above, **W** can correspond to minimum-norm estimates and their generalizations, or to stacked vectors with beamformer weights (cf. Grech et al. (2008) for a review of source analysis methods). By combining equations 1 and 2, we obtain a relationship between the ground-truth and the reconstructed source activity, which is described by the resolution matrix **K** ∈ ℝ*^N_S_^*^×^*^N_S_^* :

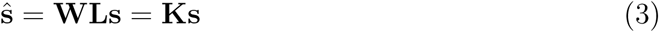

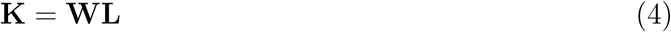

The rows of the resolution matrix **K** are referred to as the cross-talk function (CTF; Hauk et al. (2011)). CTF describes the relationship between the activity of one reconstructed and all ground-truth sources, thereby quantifying the degree of remaining field spread (RFS).

### 2.2. Extraction of ROI activity

In the present study, we apply CTF to analyze the amount and the pattern of RFS after the extraction of ROI activity. In the standard approach (see Supp. Mat. A for a review), ROI time series are obtained through aggregation of reconstructed estimates of activity for individual sources **ŝ** using a set of weights **w***_agg_* ∈ ℝ^1×^*^N_S_^* (see Table S1 for commonly used options):

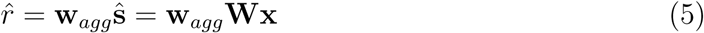

Notice that commonly used approaches can be represented by an equivalent spatial filter **w** ∈ ℝ^1×^*^N_C_^* as defined below and shown in Fig. 1A. We therefore focus on spatial filters in our discussion of different extraction pipelines:

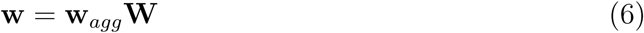

**Figure 1:**
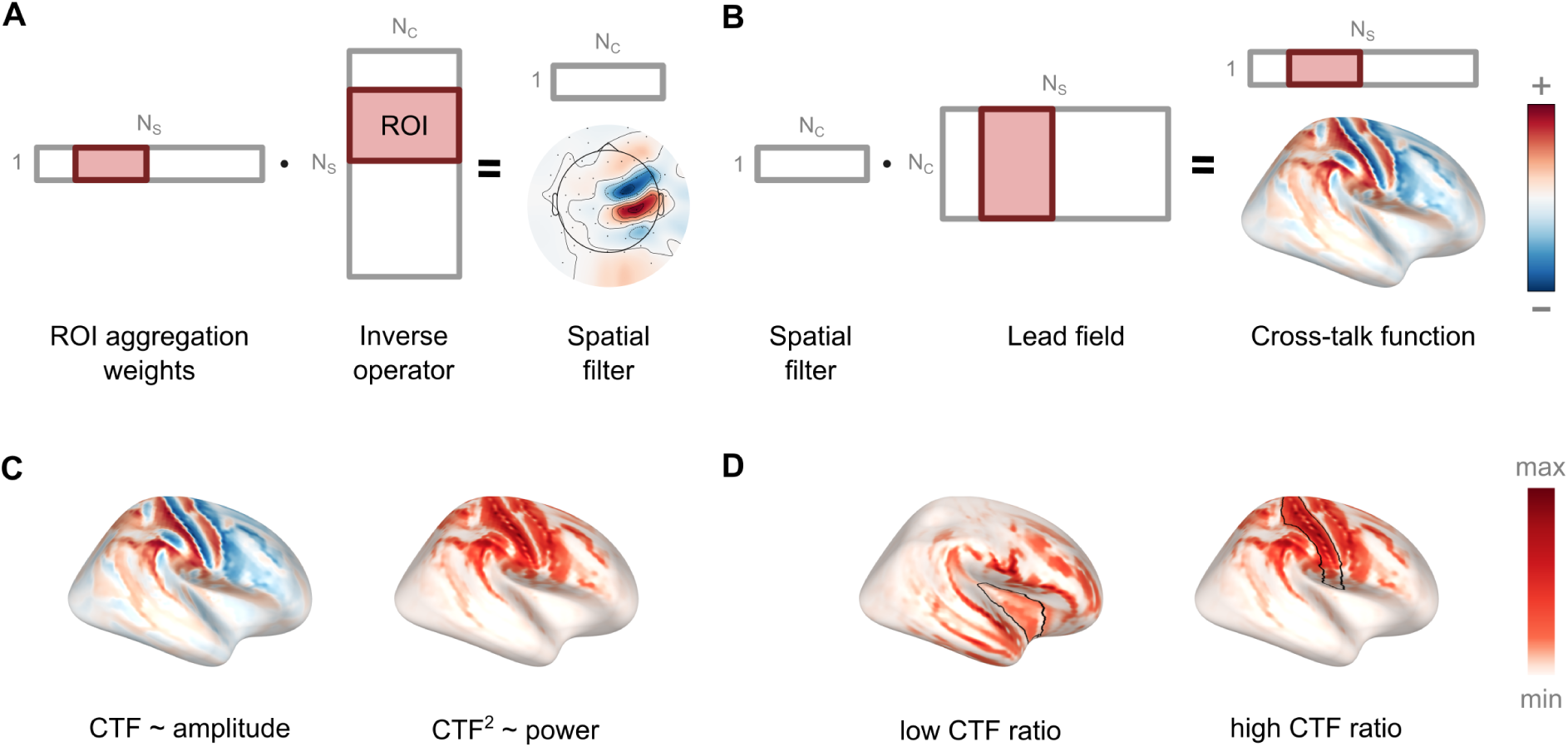
Evaluating the remaining field spread after the extraction of ROI activity using the cross-talk function (CTF). In all panels, a combination of eLORETA and mean-flip weights is used to extract ROI activity. Grey rectangles correspond to the vectors and matrices used in the derivation of CTF, and the red highlight shows the rows/columns that correspond to the sources within the target ROI. (A) An equivalent spatial filter can represent every combination of linear methods for inverse modeling and ROI aggregation. The right postcentral gyrus (Desikan-Killiany parcellation) is used as a target ROI in Panels A-C. (B) For a spatial filter, CTF shows the potential contribution of all modeled sources to the extracted time series. By default, unit variance and zero cross-covariance are assumed, but a custom source covariance matrix can be incorporated into calculations. (C) If squared, CTF reflects the potential contribution in terms of power, or the amount of variance of the ROI time series explained by each source. (D) The ratio of the norms of element-wise squared CTF within the target ROI to the whole brain can be used to evaluate the extraction of ROI activity. The higher the CTF ratio, the more likely sources within the ROI are to contribute to the extracted time series. The right insula and right postcentral gyrus are used as target ROIs to illustrate low and high CTF ratios, respectively.

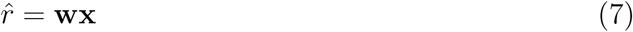

For a spatial filter **w**, it is possible to define the CTF (denoted below as **k** ∈ ℝ^1×^*^N_S_^*) in a similar way to the equation 4. The resulting CTF quantifies the relationship between the extracted time series of ROI activity *r̂* and ground-truth activity of all sources **s**:

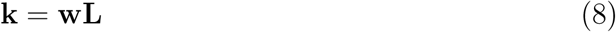

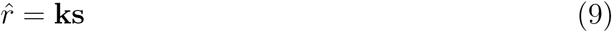

The CTF can also be visualized as a brain map, providing an intuitive way to estimate the potential main contributors of source activity (Fig. 1B). However, it is crucial to keep in mind that CTF reflects only the potential contribution of sources. In contrast, their actual contribution also depends on the covariance matrix of the source activity (Lütkenhöner and Grade de Peralta Menendez, 1997).

### 2.3. CTF reflects power contributions to the extracted time series

If we assume that the ground-truth source activity has zero mean (e.g., due to bandpass filtering) and a diagonal covariance matrix **Σ***_s_* ∈ ℝ*^N_S_^*^×^*^N_S_^*, then the power of the extracted ROI time series *P_r̂_* is a weighted sum of the ground-truth power of all sources *P_i_* = Var(*s_i_*), with weights being equal to the squared CTF elements *k_i_*^2^ (Fig. 1C):

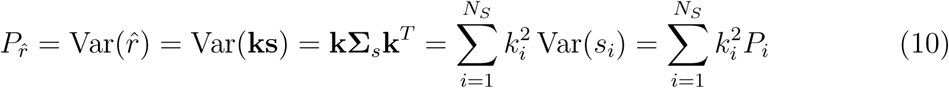

Equation 10 also shows how one can incorporate the prior knowledge in the form of the source covariance matrix in power calculations.

### 2.4. CTF ratio as a metric of extraction quality

In practice, the exact locations of ground-truth sources are often unknown. However, it is possible to evaluate the potential average contribution of sources within the ROI to the extracted signal for a given spatial filter **w** with the following ratio (later referred to as CTF ratio):

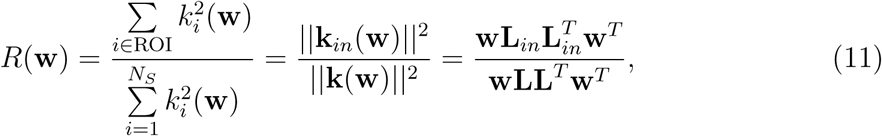

where **k***_in_* = [*k_i_* : *i* ∈ ROI] ∈ ℝ^1×^*^Nin^* is a subvector of **k** that contains only elements corresponding to sources within the ROI, **L***_in_* = [*L_ij_* : *j* ∈ ROI] ∈ ℝ*^N_C_^* ^×^*^Nin^* – a submatrix of **L** that contains only columns corresponding to sources from the ROI, *N_in_* – the number of sources in the ROI.

As shown in Supp. Mat. B.2, the CTF ratio corresponds to the expected correlation between extracted ROI time series and ground-truth activity of sources within the ROI. With a higher CTF ratio, it is more likely that sources within the target ROI will become the main contributors to the extracted time series (Fig. 1D). In Supp. Mat. B.3 we also discuss alternative definitions of the CTF ratio for arbitrary source and noise covariance matrices.

CTF ratio and its variations were previously considered by Gross and Ioannides (1999), Bolton et al. (1999), and Grosse-Wentrup et al. (2009), with all studies focusing mainly on spatial filters **w**\* that optimize the CTF ratio and can be obtained via a generalized eigenvalue decomposition:

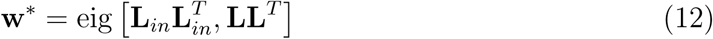

However, such optimization also defines the theoretical upper limit of the CTF ratio for a given leadfield, sensor setup, and parcellation. In the current study, we investigate the upper limit of the CTF ratio in detail and highlight the role of CTF as a diagnostic tool for all spatial filters, not only the optimal ones.

### 2.5. Effect of RFS on the estimation of inter-regional connectivity

Finally, we show how CTF can be used to investigate the effect of RFS on the estimates of functional connectivity between ROIs. Specifically, we show that the amount of spurious coherence (SC) and the effect of the RFS on the imaginary part of coherency (ImCoh; Nolte et al. (2004)) can be evaluated using CTF.

Due to RFS, the same sources may contribute to the time series of different ROIs, thereby leading to a spurious (i.e., not driven by any genuine interaction) increase in the absolute part of coherency. With CTF, we can estimate the expected amount of SC, assuming no genuine ground-truth interactions among all sources. The expected amount of SC for any pair of ROIs (denoted as *i* and *j* below) and any extraction method is equal to the dot product of normalized CTFs **k̃***_i_* and **k̃***_j_* (see Supp. Mat. B.4 for details):

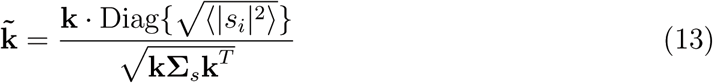

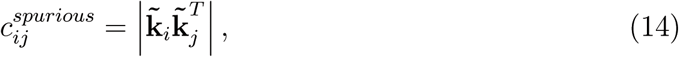

where 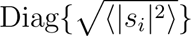 is a diagonal matrix containing the standard deviation of source activity for each source. The resulting equation for SC suggests that it depends on source amplitudes and the spatial overlap of the CTFs of spatial filters for ROIs *i* and *j*. Similar observations were made by Altukhov et al. (2023), leading to the derivation of spatial filters that minimize the spatial overlap of CTFs.

In addition, it is possible to show that the estimated ImCoh between ROIs *i* and *j* can be represented as a weighted sum of ground-truth ImCoh between all pairs of sources, with mixing weights (denoted as 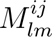 for sources *l* and *m*) depending on the normalized CTFs of both ROIs (details can be found in Supp. Mat. B.4):

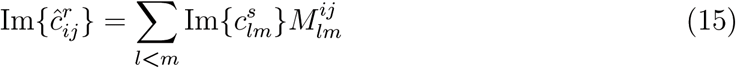

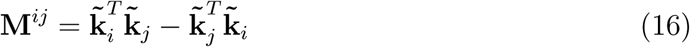

Equations 15 and 16 represent a generalization of Eq. 12 from (Sekihara et al., 2011) for the case of multiple ground-truth interactions.

## 3. Methods

### 3.1. Overview of the analyses

First, we performed two simulation experiments to illustrate the effects of remaining field spread (RFS) on the extraction of ROI activity (Experiment A) and the estimation of inter-regional connectivity (Experiment B). We then calculated the theoretical limits of the CTF ratio for commonly used parcellations and the CTF ratios achieved by several data-independent extraction pipelines. We also investigated the relationship between the theoretical limits of the CTF ratio, ROI features, and the number of recording sensors.

To validate the link between the CTF ratio and the extraction of ROI activity, we simulated 61-channel EEG data with known ground-truth source activity and conducted several experiments (Experiments 1-4b) to test the effects of deviations from the default assumptions of unit variance and zero covariance between all sources. We used real EEG data from the LEMON dataset (Babayan et al., 2019) to infer plausible values for the simulation parameters (e.g., the signal-to-noise ratio of oscillatory activity).

Finally, we used CTF to explain the consequences of RFS in real data: the level of spurious coherence derived from the resting-state EEG data of the LEMON dataset (Babayan et al., 2019) and ghost interactions using a publicly available MEG dataset from a rapid invisible frequency tagging (RIFT) experiment (later referred to as RIFT dataset; Spaak et al. (2024a)).

The analysis was mainly performed using Python 3.12.11 and the MNE-Python toolbox (v1.9.0; Gramfort et al. (2013); Larson et al. (2024)). The preprocessed data from the RIFT dataset were exported to Python using MATLAB R2023b (The MathWorks, USA) and the FieldTrip toolbox (v20241025; Oostenveld et al. (2011)). The analysis workflow was described as a set of Snakemake rules (Mölder et al., 2025).

### 3.2. M/EEG recordings and preprocessing

#### Dataset 1 (LEMON)

The LEMON dataset (Babayan et al., 2019) contains resting-state EEG recordings from 216 participants during 16 interleaved 1-minute blocks of the eyesclosed (EC) and the eyes-open (EO) conditions. Each condition lasted for 8 minutes in total. The EEG data were recorded with a BrainAmp MR plus amplifier using 61 channels placed according to the 10-10 system (locations are shown in Fig. 2A). For most steps, we used preprocessed data, which were available for 203 of 216 participants. Additionally, we used the raw data from the same participants to infer the level of amplifier noise. Three participants were excluded: two due to a mismatch in sampling frequency in the preprocessed data, and one due to missing raw data. The final sample included 200 participants (127 male; 73 female; 137: 20–40 years old; 63: 55–80 years old).

**Figure 2:**
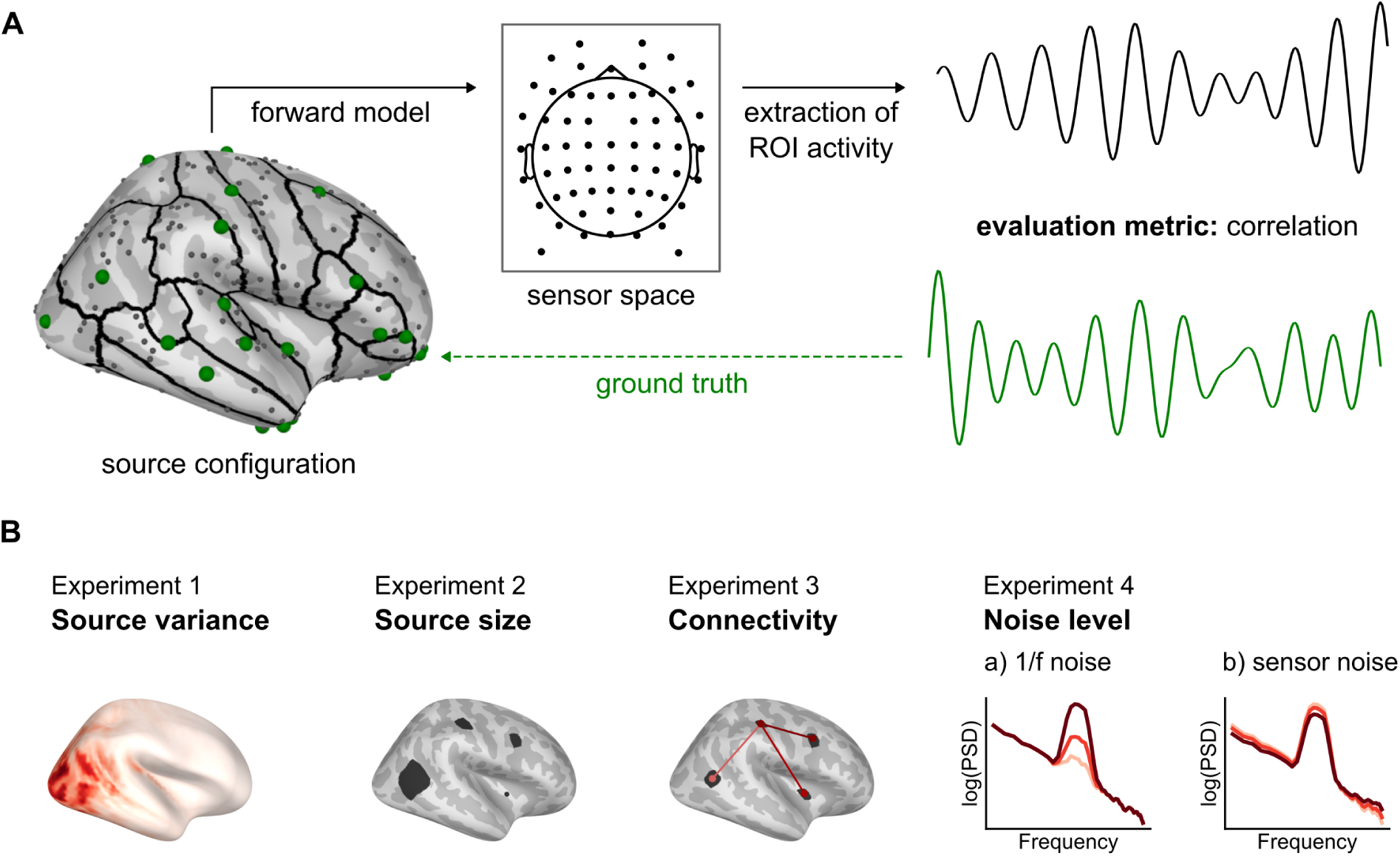
Simulations were performed to validate the CTF-based evaluation under deviation from assumptions. (A) The common workflow for the simulations. Sources of 1/f noise (dark gray) and ground-truth alpha activity (green) were simulated, and their activity was projected to the sensor space with a specified alpha-to-1/f ratio and sensor noise. The extraction quality for all pipelines was evaluated via correlation between the extracted ROI time series and the ground-truth activity waveform of the alpha source within the ROI. Both time series were filtered in 8–12 Hz before calculating the correlation. (B) We performed five experiments to test the effect of various violations of default assumptions about source activity. Specifically, we varied the spatial distribution of source variance, the spatial extent of the simulated sources, the presence and strength of ground-truth connectivity, and the level of 1/f and sensor noise. Details about the simulation parameters are presented in section 3.8. Darker shades of red denote higher variance, coherence, and noise level in Experiments 1, 3, and 4, respectively.

Preprocessing was performed by the dataset authors and included band-pass filtering between 1 and 45 Hz with an 8th-order Butterworth filter, removal of bad channels and data segments, and removal of artifactual components (i.e., eye movements/blinks, muscle activity) identified through independent component analysis (ICA). After preprocessing, the average length of the usable recordings in the EO and EC conditions was 7.8 and 7.9 minutes, respectively. For more details about the experimental setup and preprocessing, see (Babayan et al., 2019).

#### Dataset 2 (RIFT)

The RIFT dataset (Spaak et al., 2024a) contains MEG recordings of 14 participants (8 female; 6 male; mean age: 26 years, SD: 6 years) during steady-state visual stimulation at frequencies above 60 Hz. Participants completed a discrimination task and a passive viewing task, with one or two stimuli presented concurrently on the screen. In addition, the experiment included four different types of luminance modulation for the presented stimuli. In the current study, we only used the data from the passive viewing task for one and two stimuli with full amplitude luminance and contrast tagging (types 1 and 4 in the original paper; Spaak et al. (2024b)) as both resulted in high levels of coherence between the evoked steady-state responses and the presented stimuli.

MEG data were recorded using a 275-channel axial gradiometer MEG system (VSM/ CTF Systems, Coquitlam, British Columbia, Canada) in a magnetically shielded room. We used the preprocessed data, which were also available in the published dataset. Preprocessing included removal of artifactual data segments and MEG channels with excessive noise, downsampling to 600 Hz, and, finally, removal of ICA components associated with eye movements and cardiac activity. Following the authors of the dataset, we excluded two participants due to poor data quality, resulting in a final sample of 12 participants. For the analysis, we used the 0.2–1.2 s time window after the onset of the flickering stimulus. For more details about the experiment and preprocessing, see (Spaak et al., 2024b).

### 3.3. Forward model

For all analyses in source space, we used a forward model based on the fsaverage template MRI (Fischl et al., 1999) and the precomputed boundary element method solution available in MNE-Python (Gramfort et al., 2014). Unless mentioned otherwise, a source space with 4098 vertices per hemisphere (oct6 spacing in MNE-Python) was used. In simulations, we used an additional source space with 2562 vertices per hemisphere (ico4 spacing) to avoid the complete agreement between source spaces used for generating and reconstructing the simulated data, also known as the “inverse crime” (Knösche and Haueisen, 2022). The orientations of sources were fixed along the normal to the cortical surface. For most theoretical calculations and simulations, we used the 61-channel layout of the LEMON dataset with standard electrode positions. To illustrate the influence of the number of sensors on the theoretical limit of the CTF ratio, we additionally used Biosemi EEG layouts with 16, 32, 64, 128, and 256 sensors. For real-data analyses, we used the channel information provided in the LEMON and RIFT datasets.

### 3.4. Parcellations

Unless otherwise specified, we used the ROIs from the Desikan-Killiany parcellation (DK; Desikan et al. (2006)). Additionally, we considered several other parcellations based on the literature review (Supp. Mat. A) and compatibility with MNE-Python: Destrieux (Destrieux et al., 2010), Schaefer (400 ROIs, Schaefer et al. (2018)), Brodmann areas (Brodmann, 1909), and Human Connectome Project (HCP, both original and combined versions; Glasser et al. (2016)) parcellations.

To illustrate the factors that influence the theoretical limit of the CTF ratio for a given ROI, we computed the average distance from the sources within the ROI to the outer skull surface as well as the area of the cortical surface that is covered by the ROI.

### 3.5. Extraction of ROI time series

In the current study, we considered the commonly used approach of applying an inverse method followed by aggregation of the reconstructed time courses of activity for vertices within the ROI. For both steps, we selected representative methods based on the aforementioned literature review (Supp. Mat. A). Theoretical calculations were performed only for data-independent pipelines, while all pipelines were applied to the simulated data.

#### Inverse modeling

We applied eLORETA (Pascual-Marqui, 2007) with regularization parameter (*α* in the notation of the original paper) set to 0.05 and an identity noise covariance matrix. In addition, we used the LCMV beamformer with unit-noise-gain normalization (Van Veen et al., 1997; Sekihara and Nagarajan, 2008) and the same values of regularization parameter and noise covariance matrix. The data covariance matrix for LCMV was calculated using 4-second windows with 50% overlap.

#### ROI aggregation

We considered several commonly used options for aggregation weights: averaging (mean), averaging after applying the sign flip to reduce the potential amount of activity cancellation (mean-flip), taking the time course of the vertex that is the closest to the center of gravity of the ROI (centroid), and taking the first component of the singular value decomposition (SVD).

### 3.6. Estimation of functional connectivity

We used coherency as a measure of connectivity between pairs of extracted ROI time series or, for the RIFT dataset, between the ROI time series and the external stimulus. We applied Welch’s method (Welch, 1967) with a Hann window (50% overlap) and a frequency resolution of 1 Hz to obtain the cross-spectrum of both time series, and then normalized it by their autospectra to obtain coherency. Depending on the context (see Sections 3.10 and 3.11), we used either coherence (the absolute part of coherency) or the absolute value of the imaginary part of coherency (ImCoh), and averaged both measures over the frequency range of interest. Note that ImCoh is sensitive only to interactions with a non-zero phase delay (Nolte et al., 2004), which cannot be caused by RFS. In contrast, coherence is likely to detect spurious interactions driven purely by RFS.

### 3.7. Simulations

We simulated distributed ground-truth source activity (similar to a typical restingstate recording) and focused on the oscillatory activity and connectivity in the alpha (8–12 Hz) range. The simulation workflow follows existing literature (Idaji et al., 2020; Pellegrini et al., 2023) and is schematically presented in Fig. 2A. This subsection is focused on the default simulation settings. Detailed information about the performed experiments is presented in the following subsection. The mathematical formulation of the simulation algorithm is provided in the Supp. Mat. C.1, while the implementation of all building blocks is available as a Python package MEEGsim (v0.0.2; Kapralov et al. (2025)).

#### Sources and activity waveforms

In most simulations, we placed one source of alpha (8–12 Hz) activity at a randomly selected location in each ROI of the DK parcellation. By default, we used point-like sources. To make the simulations more realistic, we also considered patch-like sources, setting identical activity waveforms for all vertices within one patch-like source. Then we added 500 point-like sources of background (1/f noise with an exponent of 1) activity at random locations across the cortex. By default, the standard deviation of activity was the same for all sources and equal to 1 nAm, but in some experiments it was adjusted to introduce unequal source variance.

#### Ground-truth connectivity

To simulate waveforms with ground-truth source connectivity, we applied the Hilbert transform to the original waveform and shifted its phase by a constant value, which defined the mean phase lag. We then mixed the waveform with white noise in different proportions to control the coherence between the original and coupled waveforms. The pairs of connected sources and the mean phase lag were selected randomly, while the number and strength of connections were varied systematically.

#### Signal-to-noise ratio (SNR)

Sensor-space variance of all alpha sources was adjusted relative to all 1/f sources to obtain a specific level of global SNR in 8–12 Hz (Haufe and Ewald, 2019). Then, alpha and 1/f activity were projected to the sensor space and combined with a specified level of sensor noise (modeled as multivariate white noise).

#### Evaluation of the extraction quality

Using simulated data, we extracted the time series of activity for each ROI and evaluated the Pearson correlation between the extracted signal and the ground-truth activity of the alpha source within the ROI. Extracted and ground-truth time series were filtered in 8–12 Hz using an 8th order Butterworth filter before estimating the correlation.

### 3.8. Experiments

We performed a series of experiments (Experiments 1-4b) to test the relevance of CTF under violations of the default assumption of an identity source covariance matrix (e.g., when source activity at different spatial locations is correlated). To disentangle possible factors, we manipulated the structure of the covariance matrix in steps, as shown in Fig. 2B. In addition, we performed two simulations to illustrate the effect of RFS on the extraction of activity (Experiment A) and estimation of connectivity (Experiment B). An overview of simulation parameters for all experiments is presented in Table 1.

**Table 1:** Overview of simulation parameters in performed experiments. Values with asterisks are explained in more detail in the Methods section. Abbreviations: EO — eyes-open condition, EC — eyes-closed condition, SNR — signal-to-noise ratio, ROI — region of interest.

| Parameter | Experiment |  |  |  |  |  |  |
| --- | --- | --- | --- | --- | --- | --- | --- |
|  | 1 | 2 | 3 | 4a | 4b | A | B |
| Sampling frequency (Hz) | 250 |  |  |  |  |  |  |
| Data length (s) | 120 |  |  |  |  |  |  |
| Number of 1/f sources | 500 |  |  |  |  |  |  |
| Alpha frequency range (Hz) | 8–12 |  |  |  |  |  |  |
| Number of simulations | 100 |  |  |  |  | 1 |  |
| Location of target sources | Random within the ROI |  |  |  |  | Custom* | Center of the ROI |
| Source variance | Equal, EO-like*, EC-like* | EO-like |  |  |  | Equal, unequal* | Equal |
| Spatial extent of sources | Point | Point, patch (2, 4, 8 cm<sup>2</sup>) | Patch (2cm<sup>2</sup>) |  |  | Point | Patch (2cm<sup>2</sup>) |
| Ground-truth connectivity | None |  | None, weak*, strong* | Weak* |  | None | None, target*, interference*, both* |
| Global SNR (dB) | 4.8 |  |  | -4.8, 0, 4.8 | 4.8 | 4.8 |  |
| Level of sensor noise (%) | 1 |  |  |  | 1, 10, 25 | 1 |  |

#### Experiments 1-4b

In Experiment 1, we used point-like sources and investigated the effect of the spatial distribution of source activity, starting from equal source variance (the default assumption) and progressing toward more realistic distributions with dominant sources in the parieto-occipital areas. The source distributions were derived from the resting-state recordings of the LEMON dataset in the eyes-open (EO) and the eyes-closed (EC) conditions (Supp. Mat. C.2).

In Experiment 2, we used an EO-like distribution of source variance and varied the spatial extent of the sources. In addition to point-like sources (the default assumption), we simulated cortical patches with areas of 2, 4, and 8 cm^2^, similar to previous studies (e.g., Hincapié et al. (2017)). Patches were grown from a random location within the corresponding ROI and constrained to remain within the ROI.

In Experiment 3, we added ground-truth connectivity between sources (2 cm^2^ patches) belonging to different ROIs. We considered two levels, referred to as weak and strong, and compared them with the case of no ground-truth connectivity (default assumption). Coherence values were drawn from a Gaussian distribution. Weak ground-truth connectivity corresponded to 10 randomly selected edges with mean coherence of 0.25 and a standard deviation (SD) of 0.1. Strong connectivity was simulated by adding 20 edges with a mean coherence of 0.75 and an SD of 0.3. The phase delays were randomly selected from a uniform distribution over [0, 2*π*].

Finally, we investigated the influence of background brain activity (global SNR; Experiment 4a) and sensor noise (Experiment 4b). Both parameters were manipulated based on the values derived from the LEMON dataset (Supp. Mat. C.2). For the global SNR (as defined in Eq. S40), we considered levels of -4.8, 0, and 4.8 dB (the latter was used in all other experiments). The amount of sensor noise (see Eq. S43) was equal to 1% (default), 10%, or 25% of the total power.

#### Experiment A (activity extraction)

We placed three point-like sources near the postcentral gyrus of the left hemisphere. The source locations were selected manually to highlight the differences between the considered approaches for ROI aggregation (mean, mean-flip, centroid). To show that CTF doesn’t account for source variance by default, we simulated two cases (equal and unequal source variance) with the same source locations. In the second case, the amplitude of one source (shown in green in Fig. 3B) was increased 3-fold.

**Figure 3:**
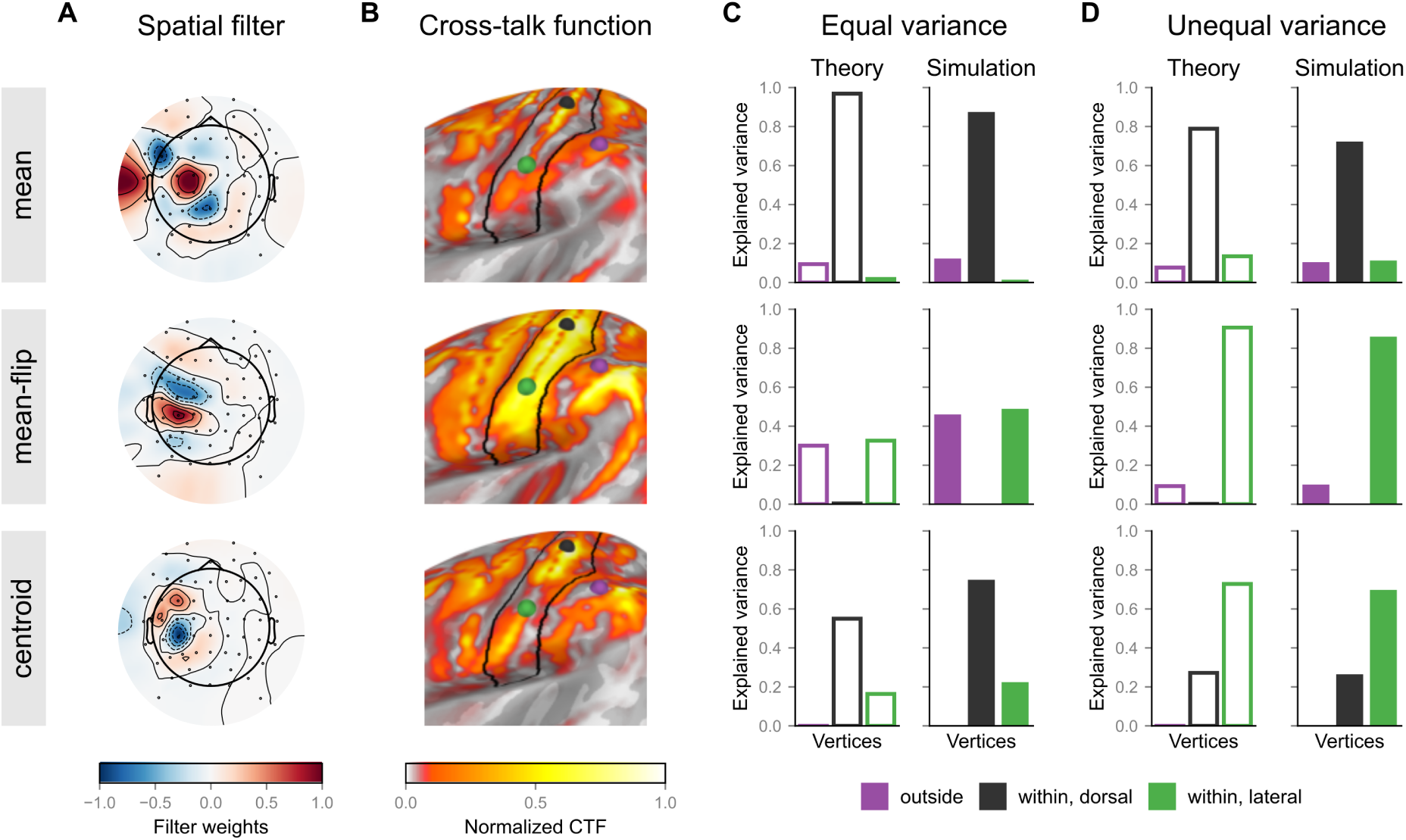
CTF can be used to investigate differences between pipelines for the extraction of ROI activity, as it shows which sources can potentially contribute to the extracted signal. Rows correspond to different aggregation methods (mean, mean-flip, centroid) after inverse modeling with eLORETA. (A) An equivalent spatial filter can represent linear pipelines for the extraction of ROI activity. (B) CTFs for each pipeline (squared element-wise). Dots show the location of sources in simulations for Panels C and D. (C) Under the assumption of equal variance, squared CTF values at the location of each simulated source correspond to the amount of variance of the ROI time series that source explains. (D) To account for unequal source variances, a source covariance matrix must be used when calculating the CTF.

#### Experiment B (connectivity estimation)

We picked two target ROIs with low CTF ratio (caudal anterior division of left cingulate cortex, isthmus division of right cingulate cortex) and two interfering ROIs (left superior frontal gyrus, right inferior parietal cortex) based on the CTF of target ROIs for eLORETA and the mean-flip aggregation approach. All ROIs were defined according to the DK parcellation. We placed a 2 cm^2^ patch-like source in the center of each ROI and varied the presence of ground-truth connections between target and interfering sources. In total, there were four possible cases: no ground-truth connections, either the target or the interfering one, or both. If a connection was present, we simulated ground-truth coherence of 0.6 and a phase lag of *π/*3.

### 3.9. CTF ratio as a metric of extraction quality

As shown in Supp. Mat. B.2, the square root of the CTF ratio is proportional to the extraction quality (correlation between ROI time series and the ground-truth activity) for a source that is placed at a random location within the ROI. Using simulated data from Experiments 1-4b, we evaluated the relationship between the CTF ratio and extraction quality using Pearson correlation and investigated the effect of deviations from the default assumptions behind CTF.

### 3.10. Spurious coherence

To estimate the amount of spurious coherence (SC) present in real resting-state EEG data, we destroyed all genuine interactions in the data with permutations (Shahbazi et al., 2010). First, the time courses of ICA components (available in the preprocessed LEMON dataset) were cropped into 2-second segments. Then, the segments were shuffled independently for each ICA component to destroy all genuine interactions present in the data. For shuffling, we used the implementation of permutations from Idaji et al. (2022). Then, the shuffled ICA time courses were projected back into the sensor space to obtain surrogate recordings. In total, we generated 100 surrogate recordings for each participant. Since all coherence that remains in the data after permutations is expected to be spurious, we estimated SC as the average coherence in the alpha (8–12 Hz) band. The values of SC were averaged across participants and surrogate recordings for the EO and EC conditions.

Since SC is directly caused by RFS, the expected amount of spurious coherence can be derived from CTFs of ROI- and pipeline-specific spatial filters (Eq. 14). We performed these calculations only for data-independent pipelines using the grand-average source variance for the respective condition (obtained with eLORETA, see Supp. Mat. C.2). For comparison, we included another model based only on the distance between the centers of gravity of the interacting ROIs. Intuitively, the amount of SC should decrease as ROIs become farther apart. However, as the precise dependency function isn’t known, we fitted a power-law model with distance between ROIs *d* as a predictor and values of SC observed in the surrogate data as the target:

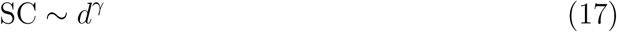

This way, we allowed the slope *γ* to be arbitrary and selected the value that yielded the best model fit.

Both models were evaluated using the Pearson correlation between their predictions and surrogate data estimates of SC. We performed two types of comparisons for each pipeline: one using raw estimates of SC (referred to as raw) and another using the difference between the pipeline-specific estimate of SC and the average estimate of SC across all pipelines (referred to as delta). The second approach assesses whether the model can explain differences in SC across extraction pipelines. For evaluation, only the upper triangular part of the SC matrices (2278 connectivity edges) was considered.

### 3.11. Analysis of RIFT data

We used the MEG data from the single-stimulus condition to estimate coherence between the time courses of activity extracted from all ROIs of the DK parcellation and the stimulus’s luminance (later referred to as brain-stimulus coherence). In the two-stimuli condition, we evaluated the absolute value of ImCoh between all pairs of ROIs (referred to as brain-brain ImCoh). In both cases, we extracted ROI time series using all data-independent pipelines as mentioned previously (Section 3.5). Connectivity values at the stimulation frequency (60 Hz) were averaged across all participants.

To illustrate the presence of brain-stimulus coherence in the data, we also estimated it in sensor space. In addition, we extracted the grand-average steady-state visual evoked response (SSVER) from the left pericalcarine cortex (DK parcellation) using eLORETA and mean-flip ROI aggregation weights.

In the main analysis, we tested whether the results of the ROI-based analyses could be explained by ghost interactions, assuming two point-like SSVER generators located in the calcarine sulci of both hemispheres. In the single-stimulus condition, we assumed that activity is identical in both hemispheres and synchronized with the external stimulus, with arbitrary coherence and phase lag. In the two-stimuli condition, we assumed a *π/*2 delay between hemispheres, since this delay was applied to the presented stimuli.

We considered three models for predicting the expected amount of brain-stimulus coherence and brain-brain ImCoh (see Supp. Mat. D for details). The first model was based on the CTF of the spatial filters that correspond to each ROI and the extraction pipeline. For comparison, we also included one model that assumed no RFS and another that assumed the RFS to depend only on the distance between the ROIs’ centroids and the assumed generators of SSVER. Since the ground-truth locations of SSVER generators were unknown, we treated them as hyperparameters for each model and performed a grid search over all locations in the pericalcarine cortex of the DK parcellation. Pearson correlation between the model predictions and estimates from real data was used as the evaluation metric. When evaluating the models, we only focused on significant connections, where the observed connectivity values exceeded the noise floor of brain-stimulus coherence or brain-brain ImCoh. The noise floor for both measures was estimated as the 95th percentile of the null distribution based on 1000 simulations with random noise of the same length as real data (see Fig. S6 and S7 for brain-stimulus coherence and brain-brain ImCoh, respectively). The final comparison between models was performed using the locations of the SSVER generators, which yielded the best fit for each model.

## 4. Results

### 4.1. CTF shows which sources potentially contribute to the extracted ROI time series

In this section, we illustrate how CTF can be used to understand the pattern of RFS for any brain area of interest and a linear ROI extraction pipeline. First, we notice that every combination of linear methods for the extraction of ROI time series can be represented by an equivalent spatial filter (Fig. 1A). For the commonly used two-step approach, the spatial filter is obtained as the product of the inverse operator matrix (e.g., eLORETA) and a vector of aggregation weights for sources within ROIs (e.g., a vector of ones for averaging). In Fig. 3A, one can clearly observe that the corresponding spatial filters differ between extraction pipelines. Still, it is unclear from the topomaps alone how these differences affect the extracted signal.

To improve the interpretability, we can multiply the spatial filter by a leadfield matrix and obtain the CTF. The CTF contains a value for each source in the forward model and describes the potential contribution of that source to the extracted ROI time series. In contrast to spatial filters, the CTFs shown in Fig. 3B provide more information and suggest that mean weights might even lead to cancellation of source activity originating from the opposite walls of the postcentral gyrus. In particular, if three sources (shown in green, black, and violet) are placed, CTF values at the source locations show the expected amount of variance that each of the sources will explain in the extracted ROI time series (Fig. 3C, theory). The predictions imply that the main contributors to the extracted signal will differ between approaches, potentially leading to differences in subsequent analyses for this configuration of sources.

However, CTF reflects only the potential impact of the source, and the amplitude of the source activity also matters in practice. If we run a simulation with equal variance of source activity for all three sources, the extraction quality obtained in the simulations will match the CTF-based predictions very closely (theory and simulation match in Fig. 3C). However, if we introduce unequal variance between the sources, for example, by increasing the amplitude of the green source by 3 times, its contribution to the extracted time series will exceed theoretical expectations (theory in Fig. 3C does not match simulations in Fig. 3D). By incorporating prior knowledge in the form of the source covariance matrix, it is possible to improve the CTF-based evaluation (theory and simulation match again in Fig. 3D).

### 4.2. CTF ratio as a metric of extraction quality: theory

In the previous section, we showed how CTF values at the exact location of the simulated sources explain which sources may contribute to the extracted ROI time series. Since the ground-truth source locations are usually unknown in resting-state data, it is more practically relevant to consider the entire region rather than focus on specific locations.

For the interpretation of the results, the main contributors to the extracted time series must be located in the considered ROI and not in other areas of the brain. Using CTF, we can assess the fidelity of extracted time series for any spatial filter by estimating how much of the total power of the extracted signal potentially originates from the ROI. We refer to this metric as the CTF ratio (Eq. 11), and examples of CTFs with low and high ratios are shown in Fig. 1D. By default, the CTF ratio is calculated assuming unit activity strength for all sources. However, any prior knowledge about the source activity can also be incorporated in the calculations.

Through optimization, it is possible to obtain a theoretical upper limit of the CTF ratio for a given lead field and ROI. Importantly, this limit cannot be exceeded by any linear spatial filter. In Fig. 4A, we show the upper limits of the CTF ratio for the DK and Schaefer parcellations. The maximal CTF ratio is much smaller in deeper regions farther from the scalp surface and thus from the recording sensors (the medial wall of both hemispheres and the insula). In addition, ROIs from the Schaefer parcellation generally exhibit a lower CTF ratio than those from the DK parcellation, in part due to differences in ROI size. These factors are illustrated for the DK parcellation in Fig. 4D and Fig. 4E, respectively. Finally, we show that the upper limit of the CTF ratio increases with the number of EEG sensors, but the increase is less pronounced for ROIs with a low CTF ratio (Fig. 4F).

**Figure 4:**
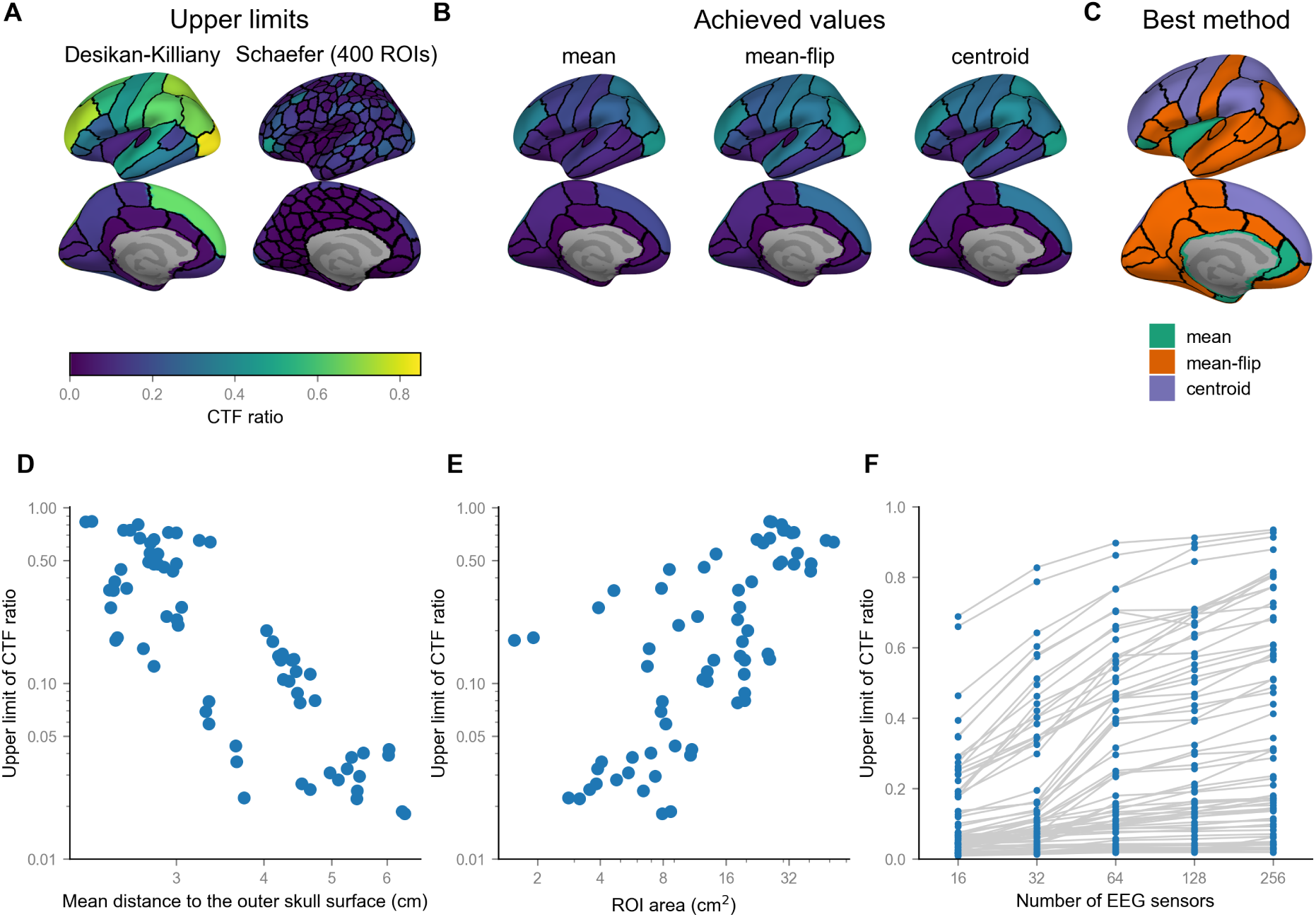
The theoretical upper limit and achieved values of the CTF ratio for the considered data-independent pipelines. (A) The upper limit of the CTF ratio for Desikan-Killiany (DK) and Schaefer parcellations. (B) Achieved values of the CTF ratio for the considered pipelines and DK parcellation. (C) The pipeline that showed the highest CTF ratio for each ROI of the DK parcellation. (D) The upper limit of the CTF ratio decreases as the average distance from sources within the ROI to the recording sensors increases. (E) The upper limit of the CTF ratio is higher for larger ROIs. (F) The upper limit of the CTF ratio increases with the number of EEG sensors. In Panels D-F, ROIs from the DK parcellation were considered.

As shown in Fig. 4B, the theoretical limits of the CTF ratio are not reached by commonly used approaches, which is at least partially caused by the applied regularization. However, the spatial distribution of the CTF ratio remains non-uniform, with deeper regions showing a lower CTF ratio. If we were to select methods that yield the highest CTF ratio across different ROIs of the DK parcellation, we would find that different methods benefit different ROIs (Fig. 4C). Therefore, the question about an optimal extraction pipeline is likely to have an ROI-specific rather than a one-size-fits-all answer.

### 4.3. CTF ratio as a metric of extraction quality: validation in simulations

We validated the idea behind the CTF ratio in simulations with distributed groundtruth source activity and used simulation parameters derived from real data, where possible. In particular, we tested whether the CTF ratio can explain differences in extraction quality between regions of interest and between approaches for the extraction of ROI time series. In addition, we empirically evaluated the reliability of CTF-based explanations when the assumption of equal activity strength for all sources does not hold.

To systemically assess the extraction quality, we simulated one ground-truth source of alpha (8–12 Hz) activity at a randomly selected location in each ROI of the DK parcellation, added 500 point-like sources of 1/f activity at random locations in the cortex, and projected the resulting source activity to the sensor space. After that, we applied several commonly used approaches for the extraction of ROI time series and, for each ROI, evaluated the correlation between the extracted signal and the ground-truth activity of the alpha source within the ROI.

By default, the CTF calculation assumes unit activity strength and zero covariance between all sources in the cortex. However, this assumption is likely violated in real M/EEG data, since, for example, sources in occipital areas typically exhibit higher alpha power in resting-state data, and connectivity can be observed even between distant regions. To test the potential value of CTF ratio in the case of violated underlying assumptions, we performed several experiments (Fig. 2B), in which we incrementally violated of the unitactivity-strength assumption by introducing a non-uniform distribution of source variance across regions, manipulating the size of alpha sources, introducing ground-truth connectivity between sources in different ROIs, and manipulating the signal-to-noise ratio in source and sensor space.

To verify that the CTF ratio predicts the differences in extraction quality between regions, we evaluated the correlation between theoretical (CTF-based) and simulation-based estimates of extraction quality across all areas of the parcellation. The results of this analysis are shown in Fig. 5A. As expected (see Fig. 3), when the CTF ratio is evaluated exactly at the ground truth location of each source with full knowledge of the source covariance matrix, the correlation with the results of simulations is almost perfect (dashed grey lines). As soon as we switch to using ROIs instead of sources, the correlation decreases slightly but remains high (solid grey lines). In addition, this correlation begins to depend on experimental conditions and worsens as the introduced deviations become stronger. A notable exception to these results is the effect of source size: as source size increases, the area of the source approaches the area of the corresponding ROI, making ROI-based and source-based evaluations more similar. Finally, when the CTF ratio is evaluated without information about the source covariance matrix and on a different source grid (solid black lines), the correlation drops to around 0.6 but remains relatively stable across all experiments.

**Figure 5:**
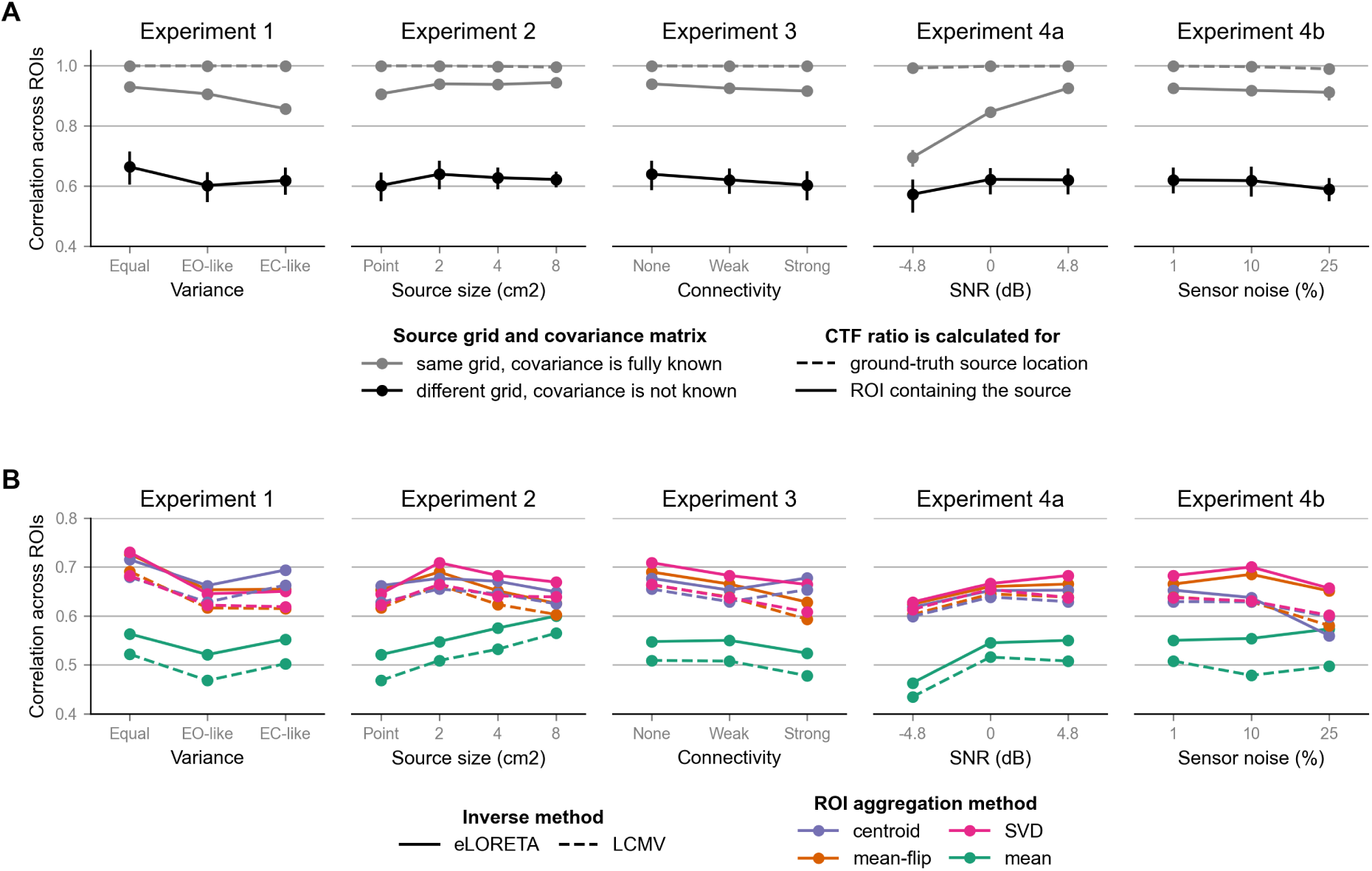
CTF ratio predicts the extraction quality (measured as the Pearson correlation between extracted ROI time series and simulated ground-truth activity) in simulations, even when the default assumption of uncorrelated unit-variance sources is violated (Experiments 1-4b). (A) Correlation between the CTF ratio and extraction quality across ROIs of the DK parcellation. The results were averaged over eight pipelines for the extraction of ROI activity. (B) Correlation between the CTF ratio and extraction quality, shown separately for each pipeline, in the case when a different source grid (compared to one used for simulating the data) and no knowledge about the source covariance matrix were used for computing the CTF (black line in Panel A). Abbreviations: EO — eyes-open condition, EC — eyes-closed condition, SNR — signal-to-noise ratio.

For the latter case, we also studied the differences between extraction pipelines (Fig. 5B). Generally, the correlation values were similar for LCMV and eLORETA. Among ROI aggregation weights, CTF-based predictions showed the lowest correlation with simulation-based estimates for averaging (mean). In addition, this method also shows different behavior in Experiment 2, in which the source size was varied. Altogether, these results might suggest a misestimation of the cancellation of activity from sources with opposite orientations when averaging is used.

### 4.4. CTF explains patterns of spurious coherence (SC) better than the distance between ROIs

From this point, we switch our focus to the estimation of inter-regional connectivity and the information that CTF can provide. One direct consequence of the remaining field spread (RFS) in the context of connectivity is the detection of spurious connectivity (e.g., as measured by coherence). When the spatial filters for two ROIs capture the same sources, high coherence may be observed even in the absence of a genuine underlying interaction. Since field spread affects commonly investigated frequencies similarly, this effect manifests as a vertical shift of the coherence spectra as the distance between the regions decreases (Fig. 6A). As CTF shows the amount of RFS, it can be used to estimate the degree of spurious coherence that would occur if no genuine ground-truth connections were present on the source level.

**Figure 6:**
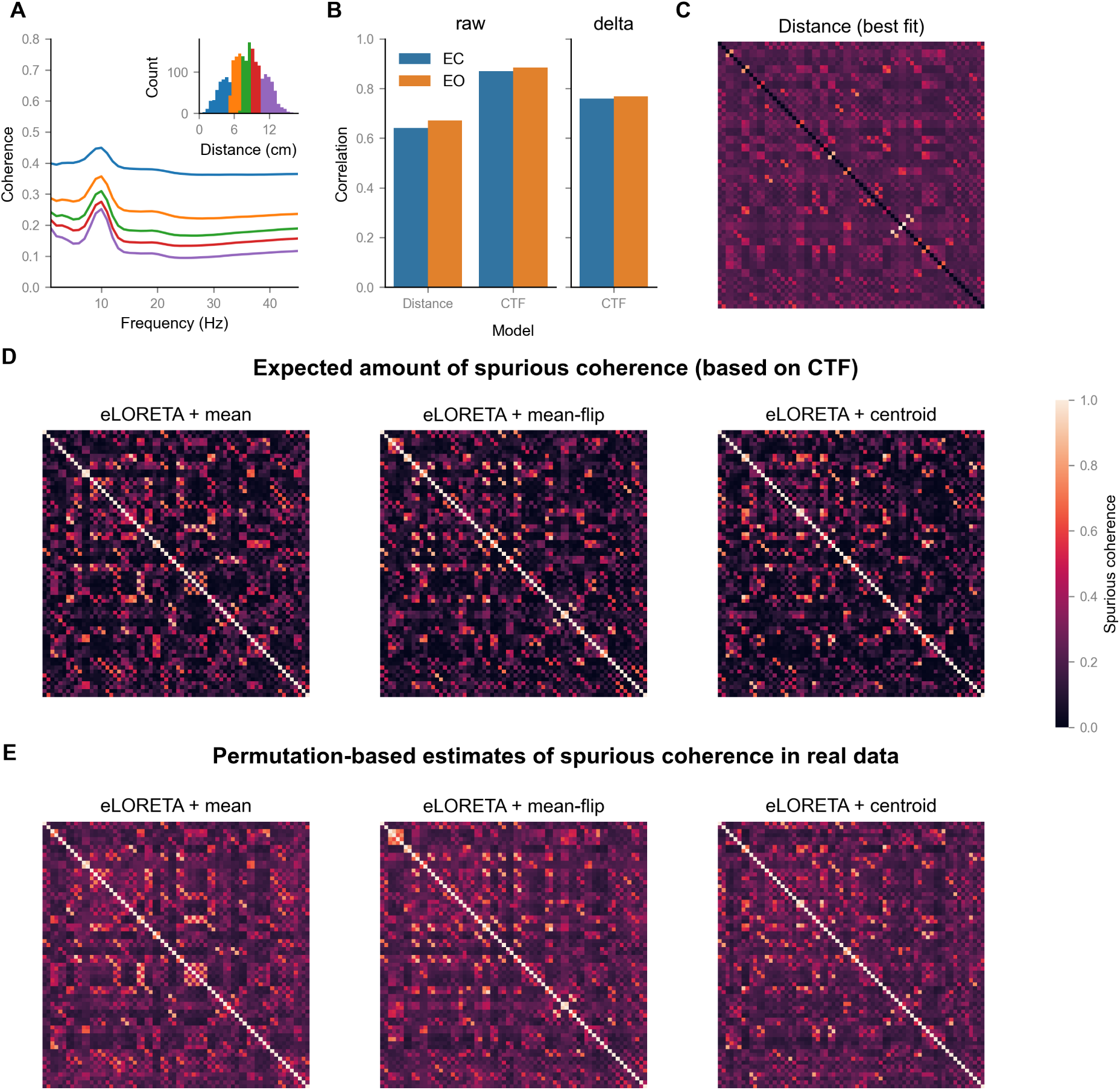
CTF predicts the degree of spurious coherence (SC) between extracted ROI time series better than the distance between ROI centroids. (A) In real data, SC manifests as a vertical shift of the grand-average coherence spectra, as shown here using resting-state data of 200 LEMON participants (eyes closed condition) for the combination of eLORETA and mean-flip. The values of SC are averaged over connectivity edges within distance-based bins (shown in the inset plot). (B) CTF explains more variance in the values of SC derived from surrogate data than the distance between ROI centers. In addition, CTF explains a considerable amount of difference in SC between the extraction pipelines. In both plots, the correlation values are averaged over three data-independent pipelines. (C) Distance-based estimates of SC for all pairs of ROIs from the DK parcellation. (D) CTF-based estimates of SC for the DK parcellation and three data-independent pipelines for the extraction of ROI activity. (E) Estimates of SC derived from surrogate data for the same parcellation and methods. In Panels C-E, ROIs are sorted alphabetically. Abbreviations: EO — eyes-open condition, EC — eyes-closed condition.

We compared the CTF-based estimates of SC (Eq. 14) with the values of SC estimated from the resting-state EEG data of the LEMON dataset using permutations. To quantify the amount of SC for each connectivity edge, we generated surrogate data using the permutation approach introduced by Shahbazi et al. (2010). This approach destroys all genuine interactions, leaving only the SC caused by the field spread. The estimates of SC for three data-independent pipelines are shown in Fig. 6E. Fig. 6D shows the CTF-based estimates of SC for the same pipelines. Additionally, we considered a competing model based on the distance between the centers of gravity of ROIs, since one might intuitively expect that SC is high between neighboring regions and decreases with distance (Fig. 6C).

We evaluated the correlation between model-based estimates of SC (based either on CTF or on the distance between ROI centroids) and estimates obtained from the surrogate data. As shown in Fig. 6B, CTF-based estimates showed a higher correlation with surrogate data than distance-based estimates (on average across considered pipelines). While the distance-based model cannot explain the difference between extraction methods by construction (distance does not depend on the extraction method), CTF also explained on average about 58% of the variance in pipeline-specific differences in SC (Fig. 6B, delta). The correlation values were similar in the eyes-closed and eyes-open conditions. The detailed results are presented in Table S3.

### 4.5. Recovery of ground-truth connectivity is not uniform across the cortex

In addition to spurious coherence, the CTF also provides information about the potential contribution of each pair of sources to the estimated inter-regional connectivity. Since extracted time series for ROIs with low CTF ratios are likely to contain activity from other ROIs, they are also more likely to capture ground-truth interactions between other brain regions. To illustrate this, we performed a simulation with a ground-truth connection between two deep ROIs with a low CTF ratio (denoted as target in Fig. 7). As shown in Fig. 7A for an exemplary extraction pipeline (eLORETA, mean-flip), the CTF for the target ROIs spreads well beyond these to other brain areas. Based on Fig. 7A, we picked two more ROIs and added another ground-truth connection between them (denoted as interference). The effect of RFS is asymmetric: the CTFs of the interfering ROIs are concentrated within these ROIs and are not (to a large extent) expected to pick up the activity of the target ones (Fig. 7C).

**Figure 7:**
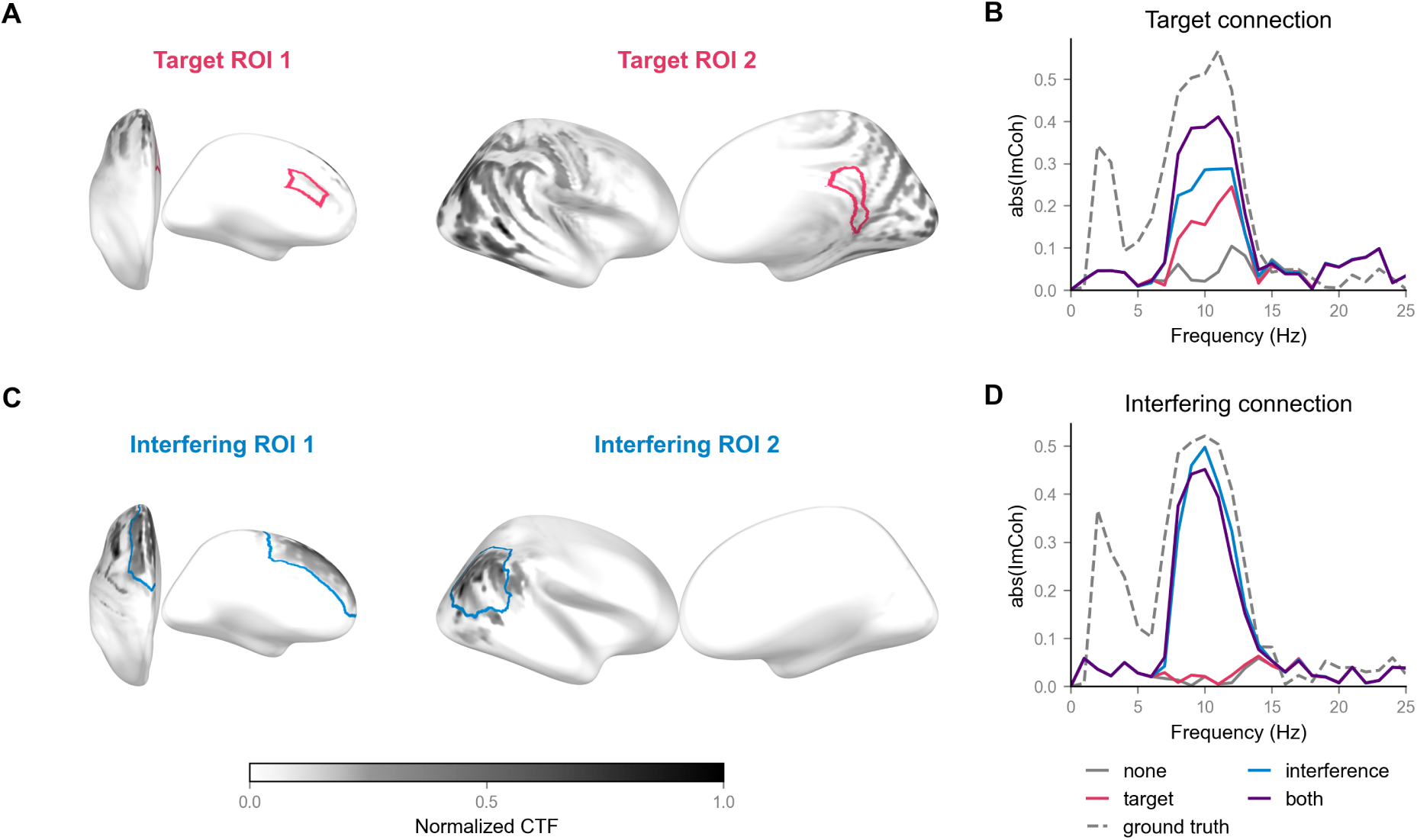
The effect of remaining field spread (RFS) on ImCoh between ROIs can be asymmetric. One target and one interference connectivity edge were simulated using ROIs from the DK parcellation. (A) Target ROIs (highlighted in red) are displayed on top of the CTF (grey), which the combination of eLORETA and mean-flip has for these ROIs. Notice that the CTF values are high within the interference ROIs (highlighted in blue in Panel C). (B) As a result, the estimated ImCoh for the target connectivity edge strongly reacts to changes in the ground-truth connectivity of the interference edge. (C) The CTF of the interference ROIs is highly localized. (D) The interference edge mostly picks up its own connectivity in simulations.

To investigate the effect of such asymmetry on the estimated connectivity values, we varied the presence of ground-truth interactions between the target and interfering ROIs by simulating configurations with no connections, one connection (either the target or the interfering one), or both. In Fig. 7B and Fig. 7D, we show the spectra of the estimated absolute ImCoh between the extracted time series of target and interfering ROIs, respectively. Notably, for both connections, ImCoh is high if and only if the interfering connection is present. As a result, both connections are dominated by ground-truth interference ImCoh, while the actual target ImCoh has a relatively small influence. This effect can be shown more generally with CTF by considering the contribution of each ground-truth ImCoh towards the estimated ImCoh between any pair of ROIs (Eq. 15). The main result remains the same: connections between ROIs with low CTF ratio are more likely to pick up interactions from the outside (Fig. S2).

### 4.6. CTF explains the connectivity patterns observed during RIFT

Finally, we tested the predictions of CTF in real data from a steady-state visual stimulation paradigm. Flickering stimuli elicit a steady-state visual evoked response (SSVER) that exhibits a strong oscillatory-like component at the stimulation frequency. Previous research (Di Russo et al., 2007; Spaak et al., 2024b) has shown that the main generators of SSVER are located in the primary visual cortex (V1), and their activity is strongly synchronized with the presented stimulus at the stimulation frequency (Fig. 8A). This phenomenon allows one to embed ground-truth connectivity in real data by controlling the amplitude and phase of luminance modulation for the presented stimuli. In sensor space, the pattern of brain-stimulus coherence in the single-stimulus condition peaks in occipital areas, as shown in Fig. 8B.

**Figure 8:**
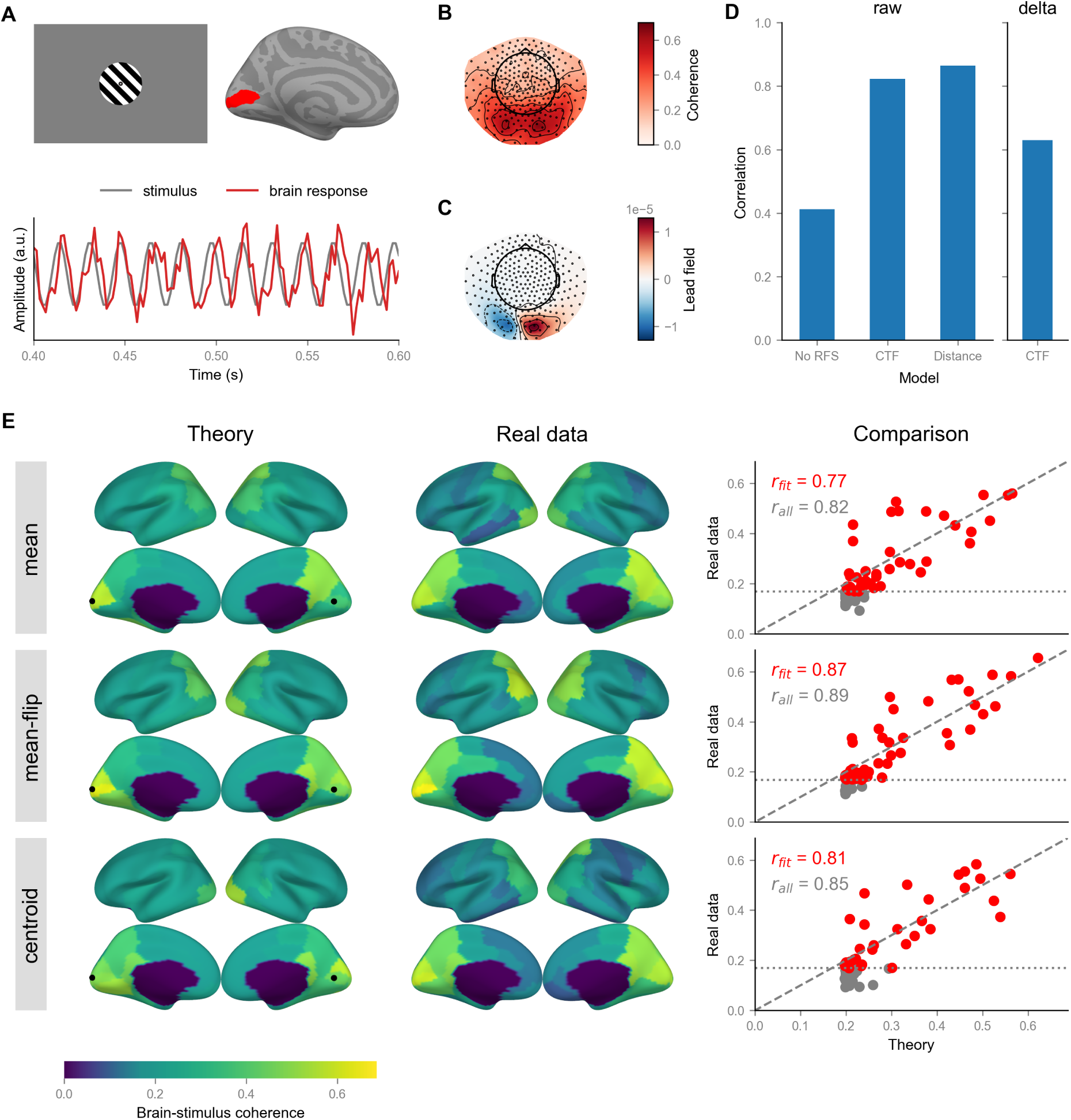
Whole-brain patterns of coherence between extracted ROI time series and the luminance of the flickering stimulus (brain-stimulus coherence) can be explained by two point-like sources located in V1 of both hemispheres using CTF. (A) When participants passively observe a flickering stimulus, a steady-state visual evoked response (SSVER) can be recorded. SSVER has a strong oscillatory-like component at the stimulation frequency (see time courses). This component is synchronized to the luminance modulation of the presented stimulus and can serve as a ground truth for connectivity benchmarks. (B) Sensor-space topomap of brain-stimulus coherence. (C) The lead field of the dipole configuration that led to the best fit for the CTF-based model. Notice its high similarity to the sensor-space pattern shown in Panel B. (D) CTF- and distance-based approaches explain a comparable amount of variance in raw values of brainstimulus coherence. Still, CTF additionally explains the difference between pipelines for ROI extraction (delta). The correlation values were computed across pooled data from three data-independent extraction pipelines. (E) Theoretical predictions based on CTF and estimates of brain-stimulus coherence from real data are strongly positively correlated for all considered pipelines. *r_fit_* and *r_all_* denote the correlation values calculated for ROIs that exceeded the noise floor (red dots) and all ROIs (gray dots), respectively. For all panels, we used the data from the experimental condition with full luminance modulation (tagging type 1) and fixed starting phase.

Due to RFS, we might observe coherence with the stimulus in other ROIs apart from V1, even if only V1 contains the main generators of SSVER. If this is the case, we should be able to predict the amount of coherence with CTF. To test this hypothesis, we used CTF to predict the expected coherence between each ROI and the external stimulus (assuming two point-like SSVER generators placed in V1) and compared the predictions with the results obtained from real data. For comparison, we considered a model with no RFS and another one in which RFS depends only on the distance between ROIs.

We evaluated the model fit only for connections that exceeded the noise floor (shown in Fig. S6 and S7 for brain-stimulus coherence and brain-brain ImCoh, respectively). The results of model comparison for brain-stimulus coherence are shown in Fig. 8D. Both CTF-based and distance-based models predict brain-stimulus coherence values better than the model without RFS. However, only the CTF-based model additionally explained 36–51% of variance between extraction pipelines in estimated brain-stimulus coherence (see Tab. S4 for detailed results for all conditions). Predictions of the CTF-based model in comparison with real data estimates are shown in Fig. 8E. Fig. 8C shows the leadfield of SSVER generators, which resulted in the best model fit. The leadfield closely matches the sensor-space topomap of brain-stimulus coherence (Fig. 8B). Similar results with slightly lower correlations were obtained for the brain-brain ImCoh in the two-stimuli condition (Fig. S8, Tab. S5).

## 5. Discussion

In this study, we applied the cross-talk function (CTF) to estimate and illustrate the effects of remaining field spread (RFS) on the extraction of M/EEG activity from regions of interest (ROIs) and on the estimation of inter-regional connectivity. We first showed that CTF can be used to evaluate potential contributions of all sources to the extracted ROI time series. Then, we used the CTF ratio to assess the fidelity of the extracted ROI time series and to derive its ROI-specific theoretical upper limit for a given lead field and parcellation. The analysis of the upper limit of the CTF ratio across ROIs highlighted the non-uniform nature of the RFS, with lower CTF ratios and thus poorer extraction of activity in regions farther from the recording sensors. These results were validated in simulations and complemented by analyses of inter-regional connectivity in real data. Altogether, the results suggest that CTF can serve as a diagnostic tool for evaluating and comparing pipelines for the extraction of ROI activity.

However, it is important to note that CTF reflects only the potential contributions of sources, while their actual contributions also depend on the source covariance matrix (Lütkenhöner and Grade de Peralta Menendez, 1997). The majority of the presented analyses assumed distributed source activity with limited information about the groundtruth source locations, which is typical of resting-state data. Therefore, conclusions may differ for source configurations with few focal sources, e.g., corresponding to different evoked responses. In that sense, CTF remains close to a spatial filter defined in source space. Multiplying it by the source covariance matrix yields a source-space spatial pattern that can be safely interpreted (Haufe et al., 2014).

### 5.1. Non-uniformity of the RFS across the cortex

The analysis of the upper limit of the CTF ratio across ROIs highlights the nonuniform nature of RFS. Regions farther from the recording sensors generally exhibit a lower CTF ratio and, hence, poorer fidelity of the extracted time series. Similar effects are observed for the estimation of inter-regional connectivity, where regions with a low CTF ratio were more likely to display ghost interactions. These results imply that connectivity edges in ‘all-to-all’ analyses (Palva and Palva, 2012) are not represented uniformly, which should be taken into account when interpreting the results.

The observed pattern of the CTF ratio across the cortex can be explained by the overlap of the lead fields of sources from different ROIs (Krishnaswamy et al., 2017). For example, suppose we try to extract activity from the insula. In that case, our spatial filter will likely also capture the activity of sources in the temporal lobe with a similar lead field. Assuming similar source strengths in both regions, activity in the temporal cortex will always contribute more to the extracted signal, resulting in a low CTF ratio for the insula. In source localization, this effect is mitigated by re-weighting the sources according to the distance to the sensors (Lin et al., 2006). However, such an approach would not help when extracting time series of activity, since weights (i.e., a spatial filter) can only be applied to different channels. Unless the spatial filter is designed to suppress the activity of specific sources, they will continue to contribute to the extracted time series (Hauk et al., 2011; de Cheveigné, 2025).

Due to the significant variability in upper limits of CTF ratio across the cortex (Fig. 4A), it is crucial to take the non-uniformity of RFS into account when interpreting the results of ROI-based analyses. As an extreme approach, regions with a low CTF ratio can be excluded from the analysis. Otherwise, the findings for such ROIs need to be interpreted with caution and thoroughly checked for potential RFS from other ROIs. If possible, one can design experimental paradigms so that the expected contribution of interfering sources is negligible. Such approaches would ensure that variations in measured parameters within low ratio areas are not merely a reflection of corresponding differences in areas with high CTF ratios.

### 5.2. Optimizing parcellations for M/EEG analysis

The theoretical upper limit of the CTF ratio was positively correlated with the size of the ROI (Fig. 4E). This result implies that even if the researchers choose a parcellation with smaller ROIs, the effective ROI size (in terms of separation of activity from other areas) remains high for a given sensor configuration. With more sensors, the upper limit of the CTF ratio increased, but the increase was not uniform across brain areas. In addition, the effect saturated as the number of sensors increased, suggesting that the spatial density of sensors is also a critical constraint (Srinivasan et al., 1998).

Recent studies have also optimized existing parcellations for use with M/EEG, acknowledging that ROI definitions based on anatomy or cellular architecture might not be optimal in the context of field spread (Farahibozorg et al., 2018; Tait et al., 2021; Sommariva et al., 2025). These studies also used CTF and its derivatives to generate parcellations and/or evaluate the performance of specific extraction pipelines for ROIs from different parcellations. In that regard, the ROI-specific theoretical limit of the CTF ratio might provide a pipeline-agnostic measure for comparing parcellations and help identify which areas benefit most from modified definitions.

### 5.3. CTF as a diagnostic tool for extraction pipelines

Recently, the question of which of the many existing pipelines for extracting ROI activity to use has become more relevant. This variability in extraction pipelines was shown to affect the results of real-data analyses (Mahjoory et al., 2017; Brkić et al., 2023; Kapralov et al., 2024). Simulation studies (Pellegrini et al., 2023; Brkić et al., 2023) provide an opportunity to compare pipelines objectively, but the results may depend on the specifics of the simulations. In this context, CTF in general and the results presented in this study in particular may help better understand the differences between RFS patterns across extraction pipelines and in choosing a suitable pipeline for the planned analysis. As shown in simulations, CTF-based estimates of extraction quality depend on a good prior for the source covariance matrix. Without it, CTF estimates may be less precise for a particular analysis, but should still provide meaningful information on average.

In addition, we showed that commonly used methods do not reach the theoretical upper limit of the CTF ratio, at least in part due to the applied regularization. Spatial filters that optimize the CTF ratio can be obtained analytically (Gross and Ioannides, 1999; Grosse-Wentrup et al., 2009), but improvements in the CTF ratio can also result from overfitting to the head model used in the underlying calculations. The importance of this effect may depend on whether a template or an individual MRI is used to create the head model, but this requires further investigation.

### 5.4. CTF explains potential effects of RFS on connectivity in real data

Using real data, we considered two scenarios that illustrate the potential effects of RFS on connectivity – spurious and ghost interactions. The results show that both the CTF and the distance between ROIs captured general trends of spurious coherence (SC) in resting-state data and ghost interactions in the RIFT paradigm. However, only the CTF could additionally explain the differences between the extraction pipelines. In addition, a high amount of RFS may come even from distant areas (Fig. 7), suggesting that CTF might provide a better explanation of the RFS than the distance between ROIs. Below, we discuss potential applications of the CTF in connectivity analyses. While we mainly focused on coherency in the analytical derivations, the effects of RFS should be similar for other commonly used connectivity measures (Nolte et al., 2020).

A well-known approach for avoiding spurious interactions is to use robust connectivity measures that are sensitive only to interactions with non-zero phase lag. However, these measures inevitably also ignore all genuine zero-lag interactions. With CTF, it is possible to evaluate the expected amount of SC for each connection (Eq. 14) and decide whether non-robust measures such as coherence could also be safely used in a particular analysis. Alternatively, one can construct a null coherence matrix assuming no genuine underlying connections and compare it with or subtract it from the one obtained from real data (Palva and Palva, 2012; Westner et al., 2024). If coherence values are compared between several groups or conditions, differences in source power can also be accounted for by specifying the source covariance matrix (e.g., from a grand-average eLORETA reconstruction).

With the RIFT dataset, we observed a reasonably good fit for a CTF-based model with two sources located in V1 of both hemispheres, consistent with the previous literature on generators of SSVER (Di Russo et al., 2007). On the one hand, this result shows that CTF can be used to account for ghost interactions when analyzing real data, potentially, in combination with other methods (e.g., the hyperedge clustering method; Wang et al. (2018)). On the other hand, it suggests that steady-state responses may serve as a benchmark for connectivity methods with a relatively well-defined ground truth, which can be manipulated by changing the properties of the presented stimuli (Herrmann, 2001; Spaak et al., 2024b).

### 5.5. Limitations

The current study has several limitations. First, we restricted the scope of the performed simulations to ensure computational feasibility. For example, we focused only on the DK parcellation, assumed fixed source orientations, and performed stepwise manipulations of the source covariance matrix (Experiments 1-4b) without considering all possible combinations of the investigated factors. Second, we considered only one source per ROI. In the case of multiple sources, the aggregation problem would still need to be solved even if we had access to the ground-truth time series, so we limited the scope to the effects of RFS from other brain areas. Finally, although we used different source grids for simulation and reconstruction, both grids were constructed from the same anatomical MRI data. Therefore, the effect of the source of anatomical data (template vs. individual MRI) on the CTF ratio remains to be explored.

There are also two critical limitations of the CTF-based evaluation of ROI extraction pipelines. As already mentioned, CTF shows only the potential, not the actual, contributions of sources to the extracted signal (Lütkenhöner and Grade de Peralta Menendez, 1997). In addition, the idea of an equivalent spatial filter applies only to combinations of linear forward and inverse models. Approaches based on a non-linear transformation of the reconstructed source time series (power, connectivity) that is averaged across all vertices of the ROI are not compatible with CTF but performed well in some simulations (Brkić et al., 2023).

## 6. Conclusions

We used CTF as a diagnostic tool to evaluate the amount and pattern of RFS for spatial filters typically used to extract regional brain activity. We showed analytically how CTF is related to the fidelity of the extracted ROI time series and the consequences of RFS for the estimation of inter-regional connectivity (spurious and ghost interactions). These results were further validated in simulations and real data. The main caveat to keep in mind when working with the CTF is that it only shows the potential contributions of sources. In contrast, actual contributions also depend on the source amplitudes and covariances. In simulations, we showed that CTF predictions remain informative even when the default assumptions are violated, and that incorporating source covariance further increases their precision. Overall, we believe that CTF can provide valuable insights into the pattern of RFS across various contexts (ROIs, extraction pipelines, experimental conditions). To facilitate its usage, we provide an open-source implementation of the key ideas discussed in the manuscript as a Python package ROIextract.

## Data and code availability statement

Both datasets that were used in the current study are publicly available. The LEMON dataset (Babayan et al., 2019) can be accessed at: https://fcon_1000.projects.nitrc. org/indi/retro/MPI_LEMON.html. The RIFT dataset (Spaak et al., 2024a) is available for download from: https://doi.org/10.34973/8dx5-7e51. All analysis scripts are available at: https://github.com/ctrltz/crosstalk-evaluation. In addition, we provide two Python packages for (1) simulation of M/EEG activity (MEEGsim; https://meegsim.readthedocs.io) and (2) evaluation of spatial filters using CTF (ROIextract; https://roiextract.readthedocs.io/).

## Ethics statement

No data was collected in this study. For the LEMON dataset, data collection was conducted in accordance with the Declaration of Helsinki, and the study protocol was approved by the ethics committee of the medical faculty at the University of Leipzig (reference number 154/13-ff). For the RIFT dataset, the data collection was approved by the local ethics committee (CMO Arnhem-Nijmegen, Radboud University Medical Center) under the general ethical approval for the Donders Centre for Cognitive Neuroimaging (’Imaging Human Cognition’, CMO 2014/288).

## CREDiT authorship contribution statement

**Nikolai Kapralov:** Conceptualization, Data curation, Formal analysis, Investigation, Methodology, Project administration, Software, Validation, Visualization, Writing – original draft, Writing – review & editing. **Alina Studenova:** Conceptualization, Investigation, Formal analysis, Software, Writing – review & editing. **Rubén Eguinoa:** Formal analysis, Software, Writing – review & editing. **Guido Nolte:** Methodology, Writing – review & editing. **Stefan Haufe:** Supervision, Writing – review & editing. **Arno Villringer:** Funding acquisition, Supervision, Writing – review & editing. **Vadim Nikulin:** Conceptualization, Investigation, Methodology, Supervision, Writing – review & editing.

## Acknowledgments

This work was supported by the Max Planck Society and, in part, by the cooperation project between the Max Planck Society and the Fraunhofer Gesellschaft (grant: project NEUROHUM). The work of Rubén Eguinoa was supported by the Spanish Ministry of Research and Innovation under Grant PID2020-118829RB-I00. Data were provided (in part) by the Radboud University, Nijmegen, The Netherlands.

