## Supplementary Material for "Non-uniform effects of remaining field spread on the estimation of M/EEG activity and connectivity between regions of interest"

### A. Overview of methods for extraction of ROI activity

We performed a literature review to provide an overview of existing pipelines for extracting ROI activity. We identified[1] 263 papers published between 2015 and 2024 that performed ROI extraction via the two-step approach: inverse modeling followed by ROI aggregation. In these papers, we collected occurrences of various inverse and aggregation methods and tracked the brain parcellations used.

The results of the review are presented in Fig. S1. Both beamformers and variations of the minimum-norm estimate are used, with sLORETA (Pascual-Marqui, 2002), LCMV (Van Veen et al., 1997), and eLORETA (Pascual-Marqui, 2007) being the most frequently used approaches (Fig. S1-A). For ROI aggregation, averaging, the first principal component, and the time course of the ROI center of mass (centroid) are the most commonly used options (Fig. S1-B). Averaging with sign flip is also used, but slightly less often. When looking at the combinations of methods (i.e., pipelines) that are used (Fig. S1-D), it becomes clear that there is no consensus upon which approaches to use, and almost every combination of methods appears at least in one study. Finally, most studies use parcellations based on anatomical (DK, Desikan et al. (2006); AAL, Tzourio-Mazoyer et al. (2002); DKT, Klein and Tourville (2012); Destrieux et al. (2010)) or cytoarchitectonic (Brodmann, 1909) features (Fig. S1-C). Functional (Schaefer et al., 2018) and multimodal (HCP, Glasser et al. (2016)) parcellations are used to a lesser extent.

[1]We used the PubMed database and the following search query: (EEG OR MEG OR electroencephalograph* OR magnetoencephalograph*) AND source AND (connectivity OR coupling OR atlas OR parcellation OR ROI) AND Journal Article[Publication Type] AND English[Language]. Preprints were excluded from the analysis.

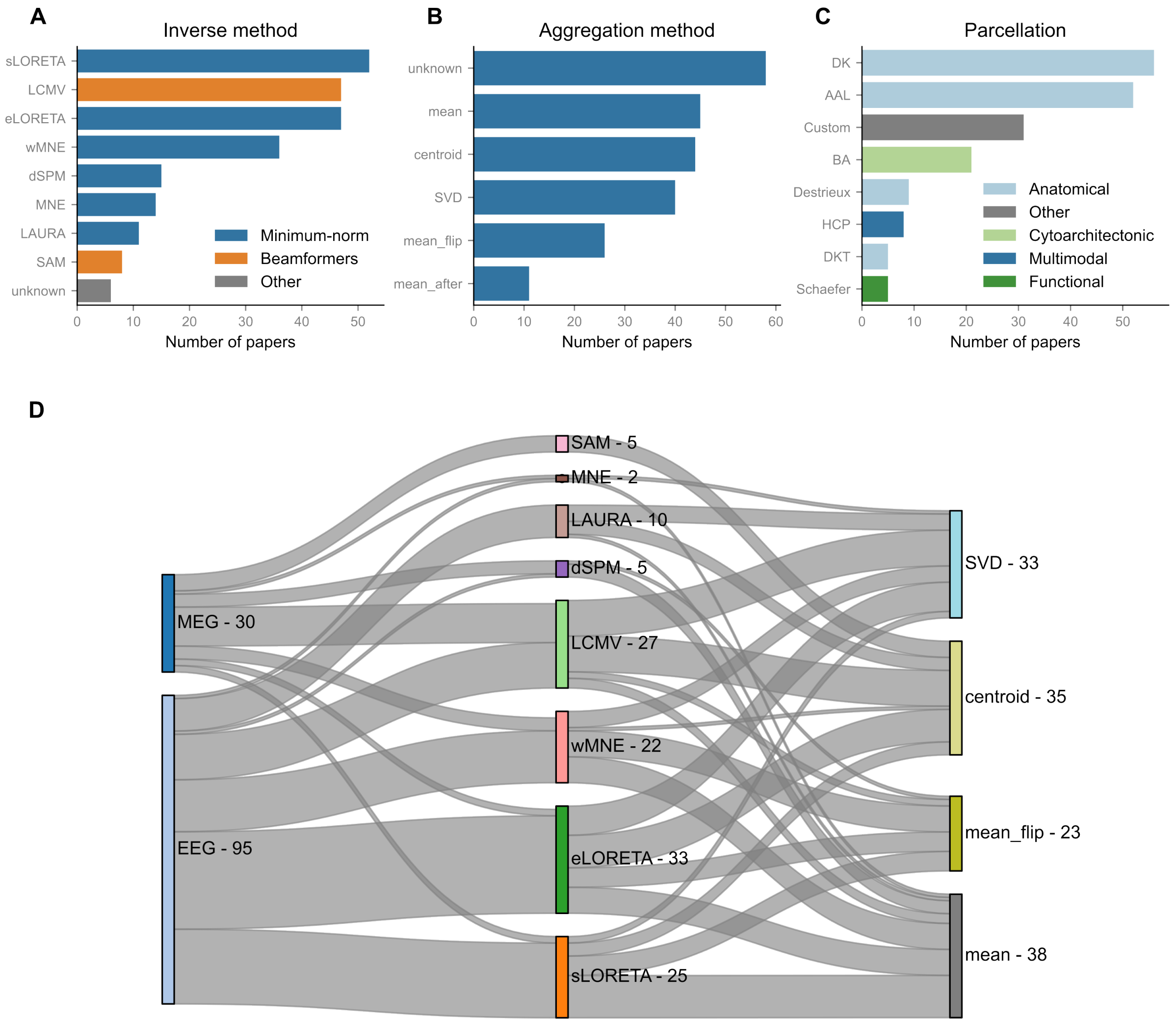


Figure S1: Literature review of pipelines for the extraction of activity from regions of interest (ROIs). Only options used by at least 5 studies are shown. (A) Inverse modeling methods used in the literature, colored by their family. (B) Approaches for ROI aggregation. See Table S1 for the detailed description. (C) Parcellations that are used to define ROIs. The parcellations are grouped by the feature types used to define them. (D) Sankey diagram of pipelines for ROI extraction shows a lack of consensus in the literature. Each pipeline connects one modality (MEG/EEG), one inverse method, and one approach for ROI aggregation. Line width denotes the number of pipelines that use each combination. Only pipelines with complete information are shown.

| Name | Weights | Explanation |
|---|---|---|
| mean | $w_i^{agg} = \begin{cases} 1, i \in I^{in} \\ 0, \text{otherwise} \end{cases}$ | Averaging reconstructed time series of all sources within the ROI |
| mean-flip | $w_i^{agg} = \begin{cases} 1 \text{ or } -1, i \in I^{in} \\ 0, \text{otherwise} \end{cases}$ | Depending on the folding of the cortex, some dipoles may have opposite orientations, thus cancelling each other's activity during averaging. With this approach, first, the dominant orientation of sources within the ROI is found via SVD. A sign flip is applied to the time series of sources whose orientation yields a negative dot product with the dominant one. After the sign flip, the time series of all the sources within the ROI are averaged. |
| centroid | $w_i^{agg} = \begin{cases} 1, i = i_c \\ 0, \text{otherwise} \end{cases}$ | Use activity from the source $i_c$, which is the closest to the ROI center of mass, to represent the whole region |
| fidelity | | Fidelity-optimized weighting operator (Korhonen et al., 2014) |
| SVD | | Weights that allow extracting the first component of the singular value decomposition of the reconstructed source time series |

Table S1: Approaches of aggregation of reconstructed time series of activity for sources within the ROI. $I^{in}$ denotes a set of indices of sources that belong to the considered ROI. Abbreviations: ROI — region of interest, SVD — singular value decomposition.

### B. Detailed theoretical derivations

#### *B.1. Notation*

Table S2 lists all variables and operators that appear in the theoretical derivations.

| Symbol | Shape | Description |
|---|---|---|
| $\mathbb{R}$ | | The set of real numbers |
| $\mathbb{E}(x)$ | | The expected value of $x$ |
| $\langle x \rangle$ | | Average value of $x$ over multiple data segments |
| $\delta_{ij}$ | | Kronecker's delta |
| $\mathbf{x}^T,\ \mathbf{X}^T$ | | Transpose of the vector $\mathbf{x}$ or matrix $\mathbf{X}$ |
| $x_i$ | | $i$-th element of the vector $\mathbf{x}$ |
| $z^*$ | | Complex conjugate of $z$ |
| $\|\|\mathbf{x}\|\|$ | | Euclidean norm of the vector $\mathbf{x}$ |
| $\mathrm{Var}(x)$ | | Variance of $x$ over time |
| $\mathrm{Cov}(x, y)$ | | Covariance of $x$ and $y$ over time |
| $\mathrm{Corr}(x, y)$ | | Correlation of $x$ and $y$ over time |
| $\mathrm{Diag}\{\mathbf{x}\}$ | | Diagonal matrix containing the elements of vector $\mathbf{x}$ |
| $\mathrm{eig}[\mathbf{A}, \mathbf{B}]$ | | Generalized eigenvalue decomposition of matrices $\mathbf{A}$ and $\mathbf{B}$ |
| $\mathrm{Im}\{z\}$ | | Imaginary part of a complex number $z$ |
| $N_C$ | | The number of M/EEG sensors |
| $N_S$ | | The number of source dipoles in the considered grid |
| $N_R$ | | The number of ROIs in the considered parcellation |
| $N_{in}$ | | The number of source dipoles in the considered ROI |
| $N_j$ | | The number of source dipoles in the $j$-th ROI |
| $\mathbf{x}(t)$ | $N_C \times 1$ | M/EEG measurements at time point $t$ |
| $\mathbf{s}(t)$ | $N_S \times 1$ | Ground-truth source activity at time point $t$ |
| $\mathbf{n}(t)$ | $N_C \times 1$ | Sensor (e.g., measurement) noise at time point $t$ |
| $\hat{\mathbf{s}}(t)$ | $N_S \times 1$ | Reconstructed source activity at time point $t$ |
| $\hat{r}(t)$ | | Extracted ROI activity at time point $t$ |
| $\mathbf{L}$ | $N_C \times N_S$ | The lead field (assuming fixed dipole orientations) |
| $\mathbf{L}_{in}$ | $N_{in} \times N_S$ | Submatrix of the leadfield that corresponds to the considered ROI |
| $\mathbf{W}$ | $N_S \times N_C$ | Inverse operator or stacked beamformer weights |

| | | |
|---|---|---|
| $\mathbf{w}_{agg}$ | $1 \times N_S$ | Weights used for aggregation of ROI activity |
| $\mathbf{w}$ | $1 \times N_C$ | Spatial filter for extraction of ROI activity |
| $\mathbf{w}^*$ | $1 \times N_C$ | Spatial filter that optimizes the CTF ratio |
| $\mathbf{K}$ | $N_S \times N_S$ | Resolution matrix |
| $\mathbf{k}(\mathbf{w})$ | $1 \times N_S$ | Cross-talk function (CTF) of the spatial filter $\mathbf{w}$ |
| $\mathbf{k}_{in}(\mathbf{w})$ | $1 \times N_{in}$ | Subvector of CTF that corresponds to the considered ROI |
| $\tilde{\mathbf{k}}(\mathbf{w})$ | $1 \times N_S$ | Normalized CTF of the spatial filter $\mathbf{w}$ |
| $R(\mathbf{w})$ | | ROI-specific CTF ratio of the spatial filter $\mathbf{w}$ |
| $P_i$, $\langle \lvert s_i \rvert^2 \rangle$ | | Power/variance of the ground-truth activity of $i$-th source |
| $P_{\hat{r}}$, $\langle \lvert \hat{r} \rvert^2 \rangle$ | | Power/variance of the extracted ROI time series |
| $\mathbf{\Sigma}_s$ | $N_S \times N_S$ | Source covariance matrix |
| $\mathbf{\Sigma}_{in}$ | $N_{in} \times N_{in}$ | Submatrix of the source covariance matrix that corresponds to the considered ROI |
| $\mathbf{\Sigma}_n$ | $N_C \times N_C$ | Noise covariance matrix |
| $\mathbf{\Sigma}_x$ | $N_C \times N_C$ | Data covariance matrix |
| $\mathbf{x}(f)$ | $N_C \times 1$ | Fourier component of M/EEG measurements at frequency $f$ |
| $\mathbf{s}(f)$ | $N_S \times 1$ | Fourier component of ground-truth source activity at frequency $f$ |
| $\hat{r}(f)$ | | Fourier component of extracted ROI activity at frequency $f$ |
| $\hat{c}^r_{ij}$ | | Estimated coherency between the time series of activity in ROIs $i$ and $j$ |
| $c^s_{ij}$ | | Ground-truth coherency between the time series of activity of sources $i$ and $j$ |
| $c^{spurious}_{ij}$ | | Spurious coherence between ROIs $i$ and $j$ |
| $\mathbf{M}^{ij}$ | $N_S \times N_S$ | Mixing weights for ImCoh |
| $\mathbf{C}$ | $N_R \times N_R \times N_R \times N_R$ | Average contribution weights in terms of ImCoh |
| $\mathbf{C}^{own}$ | $N_R \times N_R$ | Average ImCoh contribution weights for own sources |
| $\mathbf{C}^{ratio}$ | $N_R \times N_R$ | Ratio of ImCoh contribution weights for own vs. all other sources |

Table S2: Notation used throughout the theoretical derivations. The shape is only specified for vectors and matrices. Abbreviations: ROI — region of interest, ImCoh — imaginary part of coherency.

*B.2. CTF ratio corresponds to the variance explained by ground-truth sources*

In the following, we evaluate the quality of the extracted time courses of ROI activity by computing the correlation between the ground truth activity of a target source that belongs to the ROI and the ROI time series $\hat{r}$ extracted with a spatial filter $\mathbf{w}$:

$$\hat{r} = \mathbf{ks} = \sum_{i=1}^{N_S} k_i s_i \tag{S1}$$

If the activity of all sources has the same variance and no correlations ($\forall i, j\ \operatorname{Var}(s_i) = \operatorname{Var}(s_j), \operatorname{Cov}(s_i, s_j) = \delta_{ij} \operatorname{Var}(s_i)$), the CTF ratio of the target source $m$ reflects the squared correlation (i.e., explained variance) between the ROI time series and the ground-truth source activity:

$$\operatorname{Cov}(\hat{r}, s_m) = \sum_{i=1}^{N_S} k_i \cdot \operatorname{Cov}(s_i, s_m) = \sum_{i=1}^{N_S} k_i \delta_{im} \cdot \operatorname{Var}(s_m) = k_m \cdot \operatorname{Var}(s_m) \tag{S2}$$

$$\operatorname{Var}(\hat{r}) = \sum_{i=1}^{N_S} \sum_{j=1}^{N_S} k_i k_j \cdot \operatorname{Cov}(s_i, s_j) = \sum_{i=1}^{N_S} \sum_{j=1}^{N_S} k_i k_j \delta_{ij} \cdot \operatorname{Var}(s_i) = \sum_{i=1}^{N_S} k_i^2 \cdot \operatorname{Var}(s_i) \tag{S3}$$

$$\operatorname{Corr}(\hat{r}, s_m) = \frac{\operatorname{Cov}(\hat{r}, s_m)}{\sqrt{\operatorname{Var}(\hat{r}) \cdot \operatorname{Var}(s_m)}} = \frac{k_m \operatorname{Var}(s_m)}{\sqrt{\sum_{i=1}^{N_S} k_i^2 \operatorname{Var}(s_i) \cdot \operatorname{Var}(s_m)}} = \frac{k_m}{\sqrt{\sum_{i=1}^{N_S} k_i^2}} \tag{S4}$$

$$\operatorname{Corr}^2(\hat{r}, s_m) = \frac{k_m^2}{\sum_{i=1}^{N_S} k_i^2} = \frac{\mathbf{w l_m l_m}^T \mathbf{w}^T}{\mathbf{w L L}^T \mathbf{w}^T}, \tag{S5}$$

where $\mathbf{l_m}$ is the column of the lead field matrix that corresponds to the target source $m$. The final form corresponds to the CTF ratio of the target source for a given spatial filter $\mathbf{w}$. If we assume that the target source can appear in a random location within an ROI, then the expected value of the explained variance is:

$$\mathbb{E}(\operatorname{Corr}^2(\hat{r}, s_m)) = \frac{1}{N_{in}} \frac{\sum_{m \in ROI} k_m^2}{\sum_{i=1}^{N_S} k_i^2} = \frac{1}{N_{in}} \cdot \frac{\mathbf{w L}_{in} \mathbf{L}_{in}^T \mathbf{w}^T}{\mathbf{w L L}^T \mathbf{w}^T} \sim R(\mathbf{w}) \tag{S6}$$

*B.3. Variations of the CTF ratio*

In this section, we show how prior knowledge about the covariance matrices of source activity ($\mathbf{\Sigma}_s \in \mathbb{R}^{N_S \times N_S}$) and noise ($\mathbf{\Sigma}_n \in \mathbb{R}^{N_C \times N_C}$) can be taken into account in the calculation of CTF ratio. Now, we also consider sensor noise $\mathbf{n}(t) \in \mathbb{R}^{N_C \times 1}$ (time dependency is again omitted for conciseness) in the calculations:

$$\mathbf{x} = \mathbf{L}\mathbf{s} + \mathbf{n} \tag{S7}$$

The ROI time series can be extracted using a spatial filter $\mathbf{w}$:

$$\hat{r} = \mathbf{w}\mathbf{x} = \mathbf{w}\mathbf{L}\mathbf{s} + \mathbf{w}\mathbf{n} \tag{S8}$$

The total power $P_{\hat{r}}$ of the extracted ROI time series $\hat{r}(t)$ is then equal to:

$$P_{\hat{r}} = \mathbf{w}\mathbf{\Sigma}_x\mathbf{w}^T = \mathbf{w}\mathbf{L}\mathbf{\Sigma}_s\mathbf{L}^T\mathbf{w}^T + \mathbf{w}\mathbf{\Sigma}_n\mathbf{w}^T, \tag{S9}$$

where $\mathbf{\Sigma}_x \in \mathbb{R}^{N_C \times N_C}$ is the data covariance matrix. Assuming that the activity that comes from within and outside the ROI isn't correlated, the CTF ratio can be adjusted as follows:

$$R(\mathbf{w}) = \frac{\mathbf{w}\mathbf{L}_{in}\mathbf{\Sigma}_{in}\mathbf{L}_{in}^T\mathbf{w}^T}{\mathbf{w}\mathbf{L}\mathbf{\Sigma}_s\mathbf{L}^T\mathbf{w}^T + \mathbf{w}\mathbf{\Sigma}_n\mathbf{w}^T}, \tag{S10}$$

where $\mathbf{\Sigma}_{in}$ is a sub-matrix of $\mathbf{\Sigma}_s$ containing only rows and columns that correspond to sources from the target ROI. The denominator of equation S10 can also be evaluated using the data covariance matrix, making the CTF ratio data-dependent:

$$R(\mathbf{w}) = \frac{\mathbf{w}\mathbf{L}_{in}\mathbf{\Sigma}_{in}\mathbf{L}_{in}^T\mathbf{w}^T}{\mathbf{w}\mathbf{\Sigma}_x\mathbf{w}^T} \tag{S11}$$

Objective function S11 was proposed by Grosse-Wentrup et al. (2009) to serve as a backup for data-dependent spatial filters in case of low signal-to-noise ratio (SNR).

*B.4. Effect of the remaining field spread on estimation of inter-regional connectivity*

In this section, we switch to the frequency domain and, instead of the time courses $(\mathbf{x}(t), \mathbf{s}(t), \hat{r}(t))$, we consider their Fourier components at an arbitrary frequency $f$ (i.e., $\mathbf{x}(f), \mathbf{s}(f), \hat{r}(f)$). For conciseness, we omit the frequency from the following equations, since the presented equations hold for any frequency.

We can use CTF to analyze the effects of remaining field spread on the estimates of coherency between ROIs. Let $\hat{r}_i$ and $\hat{r}_j$ be the Fourier components of the time series of M/EEG activity obtained for regions $i$ and $j$ with any linear spatial filter, then:

$$\begin{aligned} \hat{r}_i &= \mathbf{k}_i\mathbf{s} = \sum_{l=1}^{N_S} k_{il}s_l \\ \hat{r}_j &= \mathbf{k}_j\mathbf{s} = \sum_{m=1}^{N_S} k_{jm}s_m \end{aligned} \tag{S12}$$

The cross-spectrum of $\hat{r}_i$ and $\hat{r}_j$ can be split into spurious (caused by the remaining field spread only) and genuine (driven by ground-truth source connectivity) parts:

$$\begin{aligned}\langle \hat{r}_i \hat{r}_j^* \rangle &= \sum_{l=1}^{N_S} \sum_{m=1}^{N_S} k_{il} \langle s_l s_m^* \rangle k_{jm} = \\ &= \underbrace{\sum_{l=m} k_{il} \langle |s_l|^2 \rangle k_{jm}}_{\text{spurious part}} + \underbrace{\sum_{l \neq m} k_{il} \langle s_l s_m^* \rangle k_{jm}}_{\text{genuine part}}\end{aligned} \tag{S13}$$

In the equation above, $\langle \cdot \rangle$ denotes averaging over data segments, which is commonly performed to get a more stable estimate of the cross-spectrum (e.g, in the Welch's method; Welch (1967)). The *spurious* part of the cross-spectrum does not depend on genuine interactions and is always present when sources contribute to the time series of both ROIs. If there are no ground-truth interactions on the source level, we can expect the following amount of spurious coherence:

$$\hat{c}_{ij}^{spurious} = \frac{\left| \langle \hat{r}_i \hat{r}_j^* \rangle^{spurious} \right|}{\sqrt{\langle |\hat{r}_i|^2 \rangle \langle |\hat{r}_j|^2 \rangle}} = \frac{\left| \sum_{l=1}^{N_S} k_{il} \langle |s_l|^2 \rangle k_{jl} \right|}{\sqrt{\left( \mathbf{k}_i \mathbf{\Sigma_s} \mathbf{k}_i^T \right) \cdot \left( \mathbf{k}_j \mathbf{\Sigma_s} \mathbf{k}_j^T \right)}}, \tag{S14}$$

If we scale the CTF values by source amplitudes and normalize using the source covariance matrix, the spurious coherence can be expressed as the dot product of normalized CTFs:

$$\tilde{\mathbf{k}} = \frac{\mathbf{k} \cdot \mathrm{Diag}\{\sqrt{\langle |s_i|^2 \rangle}\}}{\sqrt{\mathbf{k} \mathbf{\Sigma}_s \mathbf{k}^T}} \tag{S15}$$

$$\hat{c}_{ij}^{spurious} = \left| \tilde{\mathbf{k}}_i \tilde{\mathbf{k}}_j^T \right|, \tag{S16}$$

where $\mathrm{Diag}\{\sqrt{\langle |s_i|^2 \rangle}\}$ is a diagonal matrix containing the amplitude of source activity for each source. The resulting equation for spurious coherence suggests that it depends on source amplitudes and the spatial overlap of the CTFs of spatial filters for ROIs $i$ and $j$.

The *genuine* part of the cross-spectrum is present if and only if there are ground-truth interactions between sources. For the imaginary part of the cross-spectrum, which is insensitive to the spurious part, it is possible to develop the equation further:

$$\begin{aligned}\mathrm{Im}\{\langle \hat{r}_i \hat{r}_j^* \rangle\} &= \sum_{l \neq m} k_{il} \, \mathrm{Im}\{\langle s_l s_m^* \rangle\} k_{jm} = & \text{(S17)} \\ &= \sum_{l < m} (k_{il} \, \mathrm{Im}\{\langle s_l s_m^* \rangle\} k_{jm} + k_{im} \, \mathrm{Im}\{\langle s_m s_l^* \rangle\} k_{jl}) = & \text{(S18)} \\ &= \sum_{l < m} \mathrm{Im}\{\langle s_l s_m^* \rangle\} \cdot (k_{il} k_{jm} - k_{im} k_{jl}) = & \text{(S19)} \\ &= \sum_{l < m} \mathrm{Im}\{c_{lm}^s\} \cdot \sqrt{\langle |s_l|^2 \rangle \langle |s_m|^2 \rangle} \cdot (k_{il} k_{jm} - k_{im} k_{jl}) & \text{(S20)}\end{aligned}$$

To obtain coherency, the cross-spectrum is normalized by the variance of the ROI time series:

$$\mathrm{Im}\{\hat{c}^r_{ij}\} = \frac{\mathrm{Im}\{\langle \hat{r}_i \hat{r}^*_j \rangle\}}{\sqrt{\langle |\hat{r}_i|^2 \rangle \langle |\hat{r}_j|^2 \rangle}} = \sum_{l<m} \mathrm{Im}\{c^s_{lm}\} \frac{\sqrt{\langle |s_l|^2 \rangle \langle |s_m|^2 \rangle} \cdot (k_{il}k_{jm} - k_{im}k_{jl})}{\sqrt{\left(\mathbf{k}_i \mathbf{\Sigma_s} \mathbf{k}_i^T\right) \cdot \left(\mathbf{k}_j \mathbf{\Sigma_s} \mathbf{k}_j^T\right)}} \tag{S21}$$

Using the normalization introduced in Equation S15, we can extract and simplify the weights, with which the ground-truth ImCoh values between all pairs of sources get mixed into the estimated ImCoh between ROIs $i$ and $j$:

$$\mathbf{M}^{ij} = \tilde{\mathbf{k}}_i^T \tilde{\mathbf{k}}_j - \tilde{\mathbf{k}}_j^T \tilde{\mathbf{k}}_i \in \mathbb{R}^{N_S \times N_S} \tag{S22}$$

$$\mathrm{Im}\{\hat{c}^r_{ij}\} = \sum_{l<m} \mathrm{Im}\{c^s_{lm}\} M^{ij}_{lm} \tag{S23}$$

*B.5. Non-uniform effects of RFS on the estimation of ImCoh*

As shown in Eq. S22, the mixing of ImCoh is described by a separate matrix $\mathbf{M}^{ij}$ for each pair of ROIs $i$ and $j$. In this section, we analyze the average contribution weights (in terms of ImCoh) for sources from all ROIs for the Desikan-Killiany parcellation. First, we construct the contribution matrix $\mathbf{C} \in \mathbb{R}^{N_R \times N_R \times N_R \times N_R} = [C_{ijkl}]$ as follows:

$$C_{ijkl} = \frac{1}{N_k N_l} \sum_{m \in \mathrm{ROI}_k} \sum_{n \in \mathrm{ROI}_l} |M^{ij}_{mn}| \tag{S24}$$

The value $C_{ijkl}$ describes how strongly (on average) the ground-truth values of ImCoh between all pairs of sources from ROIs $k$ and $l$ will contribute to the estimated ImCoh between ROIs $i$ and $j$. Importantly, it only shows the weight of the potential contribution, so if the ImCoh between sources is zero, there will be no actual contribution from these sources. The expected contribution weight of *own* sources (the sources that belong to ROIs $i$ and $j$) is equal to:

$$C^{own}_{ij} = C_{ijij} \tag{S25}$$

Ideally, own sources should contribute as much as possible so that we can attribute the results of the connectivity analysis to the respective ROIs. As shown in Fig. S2-B for an extraction pipeline based on eLORETA and mean-flip, the contribution weights for ImCoh of own sources are not uniform across the cortex (assuming an identity source covariance matrix). ROIs with higher CTF ratio (see Fig. S2-A for the color scheme) are generally expected to get a larger contribution in terms of ImCoh from their own sources.

This effect becomes even clearer if one considers the ratio of contribution weights of own sources to the maximum contribution weights from sources that belong to all other pairs of ROIs:

$$C_{ij}^{ratio} = \frac{C_{ij}^{own}}{\max\limits_{k,l \neq i,j} C_{ijkl}} \tag{S26}$$

This ratio is shown in Fig. S2-C, implying that regions with low CTF ratio are more likely to extract ImCoh of other connections rather than their own ones ($C_{ij}^{ratio} < 1$, shown in light blue). The opposite can be expected for regions with a high CTF ratio.

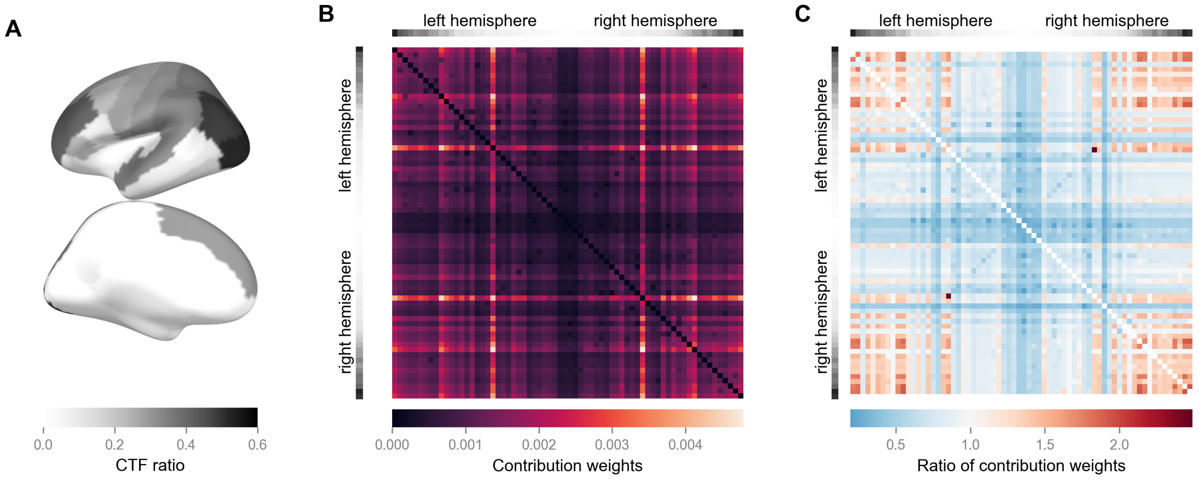


Figure S2: Recovery of ground-truth connectivity (measured with the absolute value of ImCoh) is not uniform across the cortex. The visualization of connectivity matrices is adapted from Liljeström et al. (2015). ROIs are grouped by the hemisphere they belong to and sorted by the CTF ratio within hemispheres. (A) CTF ratio achieved by the combination of eLORETA and mean-flip weights, averaged over homologous pairs of ROIs from both hemispheres. (B) Average contribution weights of ground-truth ImCoh from sources that belong to the target pair of ROIs (Eq. S25) are not uniform across the cortex. Regions that exhibit a higher CTF ratio generally have higher weights for their own ground-truth connections. (C) Deeper regions with low CTF ratio (insula, medial walls of both hemispheres) are more sensitive to ground-truth ImCoh from other connections than their own ones, as indicated by values of contribution ratio (Eq. S26) lower than 1. The opposite holds for ROIs close to the recording sensors, which have a higher CTF ratio.

### C. Detailed description of simulations

#### *C.1. Mathematical description of the simulation algorithm*

In this section, we describe the mathematical equations behind our approach for simulating EEG data. The following steps were performed:

1. *Noise sources.* We placed $N_{noise} = 500$ point-like sources at randomly selected locations within the source space. Power-law noise with the exponent of 1 was used as a waveform of activity for each noise source $\mathbf{s}_i^{noise}(t), i \in [1, N_{noise}]$. The noise was generated using the algorithm of Timmer and König (1995), as implemented in the `colorednoise` Python package.

2. *Sources of alpha activity.* We placed either point-like or patch-like sources of alpha activity in each ROI of the DK parcellation. The total number of alpha sources $N_{alpha}$ was therefore equal to the number of ROIs $N_R$. To obtain waveforms of oscillatory activity $\mathbf{s}_j^{alpha}(t)$ for each alpha source $j \in [1, N_{alpha}]$, we generated white noise and filtered it in 8-12 Hz range using an 8th order Butterworth filter. In the case of patch-like sources, the time courses of activity for all vertices belonging to one source had identical waveforms and zero delays between each other.
3. *Connectivity.* To set ground-truth connectivity between a pair of sources (denoted as 1 and 2), we used the following approach. First, a constant phase delay $\Delta\varphi$ was applied to the activity waveform of the first source $s_1^{alpha}(t)$ via the Hilbert transform. In the equations below, $A_1(t)$ and $\varphi_1(t)$ denote the instantaneous amplitude and phase, while $j$ stands for the imaginary unit:

$$s_1^{alpha}(t) \equiv \mathrm{Re}\left\{A_1(t) \cdot \exp\left(j\varphi_1(t)\right)\right\} \tag{S27}$$

$$s_1^{shift}(t) = \mathrm{Re}\left\{A_1(t) \cdot \exp\left(j\varphi_1(t) + j\Delta\varphi\right)\right\} \tag{S28}$$

To control the coherence between the sources, we mixed the shifted version of the original waveform with a specific amount of white noise $n(t)$. The signal-to-noise ratio (SNR) $\gamma_c$ in 8-12 Hz range determines the resulting coherence $c_{12}$ between the original waveform and such mixture (Bendat and Piersol, 2010):

$$\gamma_c = \frac{\mathrm{Var}_{\text{8-12 Hz}}\left(s_1^{shift}(t)\right)}{\mathrm{Var}_{\text{8-12 Hz}}\left(n(t)\right)} \tag{S29}$$

$$c_{12} = \sqrt{\frac{\gamma_c}{1+\gamma_c}} \tag{S30}$$

The required SNR can be derived from the desired coherence $c_0$ as follows:

$$\gamma_{c0} = \frac{c_0^2}{1-c_0^2} \tag{S31}$$

To obtain such an SNR, the amplitude of the shifted waveform needs to be adjusted by the following factor:

$$f_c = \sqrt{\frac{\gamma_{c0} \cdot \mathrm{Var}_{\text{8-12 Hz}}\left(n(t)\right)}{\mathrm{Var}_{\text{8-12 Hz}}\left(s_1^{shift}(t)\right)}} \tag{S32}$$

The final waveform of the second source is then defined as:

$$s_2^{alpha}(t) = f_c \cdot s_1^{shift}(t) + n(t) \tag{S33}$$

We used depth-first search to traverse the connectivity graph and ensure that all waveforms are set correctly when multiple connectivity edges need to be simulated.

4. *Normalization.* Generated noise waveforms were divided by their standard deviation (SD) over time to normalize the SD across sources:

$$\tilde{s}_i^{noise}(t) = \frac{s_i^{noise}(t)}{\sqrt{\mathrm{Var}(s_i^{noise})}} \tag{S34}$$

Waveforms of alpha activity were also normalized to have the same SD for all sources. To make the total variance of patch activity independent of the number of vertices in the patch ($N_v$), we also included the number of vertices in the normalization ($N_v = 1$ for point sources). Finally, in some experiments, we manipulated the spatial distribution of the SD of alpha activity (e.g., to mimic the dominance of parieto-occipital alpha sources, which is typical in resting-state recordings). To achieve this, the waveforms of alpha activity were scaled by the desired SD $\sigma$ that corresponded to the location of the source (or its center in case of patches). The combined normalization is therefore:

$$\tilde{s}_j^{alpha}(t) = \frac{s_j^{alpha}(t) \cdot \sigma}{\sqrt{\mathrm{Var}(s_j^{alpha}) \cdot N_v}} \tag{S35}$$

5. *Signal-to-noise ratio (SNR).* We adjusted the SNR of alpha activity based on the approach described in Haufe and Ewald (2019). First, we projected the activity of all noise and point-/patch-like sources to sensor space using the corresponding parts of the lead field matrix ($\mathbf{L}_i$ contains only columns that correspond to the vertices of the $i$-th source):

$$\mathbf{x}^{noise}(t) = \sum_{i=1}^{N_{noise}} \mathbf{L}_i \tilde{s}_i^{noise}(t) \tag{S36}$$

$$\mathbf{x}^{alpha}(t) = \sum_{j=1}^{N_{alpha}} \mathbf{L}_j \tilde{s}_j^{alpha}(t) \tag{S37}$$

Then, we calculated the mean variance of the projected noise and alpha activity across $N_C$ sensors after filtering the time series in 8-12 Hz range:

$$P_{noise} = \frac{1}{N_C} \sum_{i=1}^{N_C} \mathrm{Var}_{8\text{-}12\ \mathrm{Hz}}(x_i^{noise}(t)) \tag{S38}$$

$$P_{alpha} = \frac{1}{N_C} \sum_{j=1}^{N_C} \mathrm{Var}_{8\text{-}12\ \mathrm{Hz}}(x_j^{alpha}(t)) \tag{S39}$$

The global SNR of alpha relative to 1/f activity $\gamma_{alpha}$ was then defined as:

$$\gamma_{alpha} = \frac{P_{alpha}}{P_{noise}} \tag{S40}$$

To obtain the target SNR $\gamma_0$, we scaled the amplitude of alpha activity by the following factor:

$$f_{\text{SNR}} = \sqrt{\frac{\gamma_0 \cdot P_{noise}}{P_{alpha}}} \tag{S41}$$

6. *Projecting to sensor space.* The waveforms of alpha activity and 1/f noise were projected to sensor space and summed:

$$\mathbf{x}^{brain}(t) = f_{\text{SNR}} \cdot \sum_{j=1}^{N_{alpha}} \mathbf{L}_j \tilde{s}_j^{alpha}(t) + \sum_{i=1}^{N_{noise}} \mathbf{L}_i \tilde{s}_i^{noise}(t) \tag{S42}$$

Finally, we generated multivariate white noise $\mathbf{x}^{sensor}(t)$ to model sensor noise and added it to the projected brain activity to obtain the simulated data $\mathbf{x}(t)$:

$$\mathbf{x}(t) = \sqrt{1 - \gamma_{sensor}} \cdot \mathbf{x}^{brain}(t) + \sqrt{\gamma_{sensor}} \cdot \frac{P_{brain}}{P_{sensor}} \cdot \mathbf{x}^{sensor}(t) \tag{S43}$$

$$P_{brain} = \frac{1}{N_C} \sum_{i=1}^{N_C} \text{Var}_{\text{8-12 Hz}}(x_i^{brain}(t)) \tag{S44}$$

$$P_{sensor} = \frac{1}{N_C} \sum_{j=1}^{N_C} \text{Var}_{\text{8-12 Hz}}(x_j^{sensor}(t)) \tag{S45}$$

The parameter $\gamma_{sensor}$ controls the fraction of total power of the simulated data that is explained by sensor noise.

#### *C.2. Inferred values of simulation parameters*

We estimated the values of several simulation parameters (global SNR, spatial distribution of source power, and sensor noise level) using resting-state recordings from the LEMON dataset. This approach aimed to ensure that reasonable values were used in the experiments. The estimation approach aligns with the role of each parameter in the simulations.

Global SNR was estimated from the average power spectral density (PSD) across all channels (Welch's method, 0.5 Hz frequency resolution, 50% overlap). SNR was defined as the ratio of power in the alpha band (8–12 Hz) to the mean power in the flanking frequency bands: 5–7 and 13–15 Hz (Fig. S3-A). Fig. S3-B shows the obtained distributions of global SNR in the eyes-open (EO) and eyes-closed (EC) conditions. Median global SNR was equal to 1.3 (1.2 dB) and 3.6 (5.6 dB) for EO and EC conditions, respectively. In simulations, we used an SNR of 3 (4.8 dB) as the default. We considered two additional SNR levels (-4.8 and 0 dB) to test the effect of SNR on the relationship between CTF ratio and extraction quality.

To introduce unequal variance of the activity of simulated sources, we used the grand-average values of source-space alpha power. For each subject, the sensor-space data were filtered in 8–12 Hz using an 8th order Butterworth filter. Estimates of source power were obtained using eLORETA and normalized to remove between-subject differences in total power before averaging. The resulting power maps for EO and EC conditions are shown in Fig. S3-C. As could be expected, occipital and sensorimotor areas show the strongest activity.

Finally, we obtained two estimates for the level of sensor-space noise. The first one is based on the amplifier noise floor, which appears as a plateau in the PSD of the raw data at high frequencies (Scheer et al., 2006). Since the amplifier used for recording the LEMON data had a built-in low-pass filter at 1 kHz, we used the value of power at 800 Hz as an upper bound of the amplifier noise (Fig. S3-D). As shown in Fig. S3-E, amplifier noise never exceeded 1% of total alpha power. The second estimate of the noise level was obtained via a 5-fold spatial cross-validation (Hashemi et al., 2021). This approach allows estimating the amount of variance that cannot be explained by the forward and inverse models (in our case, eLORETA). Such residual variance might originate from sources of activity other than the brain (e.g., muscles). For cross-validation, we used the first 3 minutes of preprocessed data for each subject and varied the regularization parameter $\lambda$ from 0.0001 to 10 in 15 log-spaced steps. The lowest residual variance across all values of $\lambda$ was taken as the noise level. Fig. S3-F shows the distribution of noise level relative to the subject-specific alpha power, with 11.2% and 10.0% as median values for EO and EC conditions, respectively. In simulations, we set the sensor noise power to 1%, 10%, and 25% of the total power to test its effect on the relationship between the CTF ratio and extraction quality.

### D. Models of the remaining field spread for RIFT data

For all models, we assumed two dipolar generators of SSVER in V1 (pericalcarine cortex, DK parcellation). In the single-stimulus condition, we assumed that the generators have identical ground-truth time courses of activity and measured coherence between the extracted ROI time series and the external stimulus (brain-stimulus coherence). In the two-stimuli condition, we assumed a phase lag of $\pi/2$ between the ground-truth time courses since such a delay was applied to the presented stimuli, and we measured the absolute value of ImCoh between all pairs of ROIs (brain-brain ImCoh). Fig. S4-A and S4-B show the predictions of all considered models for brain-stimulus coherence and brain-brain ImCoh, respectively.

#### *D.1. No RFS*

If we assume that no RFS is present after the extraction of the ROI time series, then only the ROIs that contain SSVER generators would show non-zero coherence with the external stimulus in the single-stimulus condition and ImCoh in the two-stimuli condition.

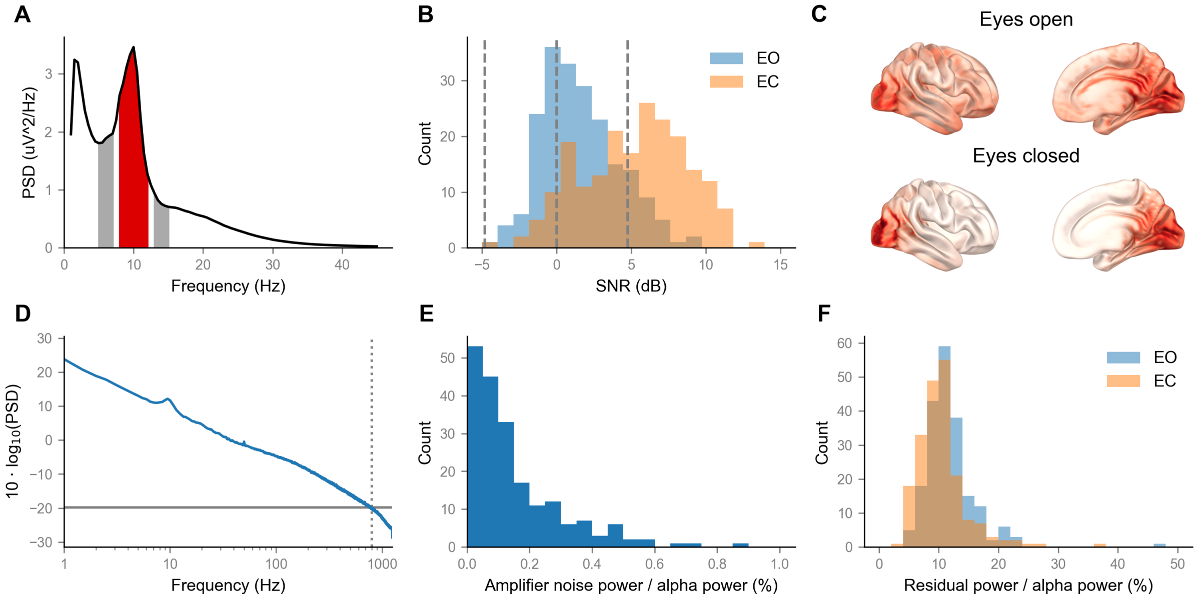


Figure S3: Simulation parameters inferred using the resting-state data from the LEMON dataset. (A) Global SNR was estimated as the ratio of power in the alpha band (red) to the mean power in the flanking frequency bands (gray). (B) Histograms of global SNR values in eyes-open (EO) and eyes-closed (EC) conditions. Dashed lines show the values of global SNR that were used in simulations. (C) Spatial distribution of the grand-average source-space alpha power. (D) Power at 800 Hz was used as the upper bound of amplifier noise. (E) The power of amplifier noise did not exceed 1% relative to the total alpha power for any participant. (F) Estimates of noise level based on spatial cross-validation in EO and EC conditions.

*D.2. Distance-based RFS*

Alternatively, we can account for the distance between ROIs and SSVER generators when evaluating the potential effects of RFS. For the $i$-th ROI, let $d_{1i}$ and $d_{2i}$ be the average Euclidean distance between the vertices of the ROI and two SSVER generators, respectively. Then, brain-stimulus coherence should be related to the average distance $d$ to both SSVER generators from that ROI:

$$d = \frac{1}{2} \cdot (d_{1i} + d_{2i}) \tag{S46}$$

In case of brain-brain ImCoh between ROIs $i$ and $j$, we used the minimum distance across pairwise combinations listed below:

$$d = \frac{1}{2} \cdot \min\left[d_{1i} + d_{2j}, d_{1j} + d_{2i}\right] \tag{S47}$$

Since the precise functional dependency of RFS on distance is not known, we fit a power-law model to the values $c_{real}$ obtained in real data with distance $d$ as a predictor, thus allowing for an arbitrary slope $\gamma$:

$$c_{real} \sim d^{\gamma} \tag{S48}$$

In Fig. S4, the predictions of the distance-based model are shown with the slope of $-1$ for illustration purposes.

*D.3. CTF-based RFS*

With CTF, we can derive the expected amount of brain-stimulus coherence and brain-brain ImCoh more precisely and additionally capture differences between pipelines for ROI activity extraction. The following holds for the Fourier components of the extracted signal $\hat{r}$ and ground-truth source activity $s_i$ at the stimulation frequency:

$$\hat{r} = \mathbf{ks} = \sum_{i=1}^{N_S} k_i s_i \tag{S49}$$

In what follows, we refer to the sources assumed to be the SSVER generators by indices 1 and 2, and denote the Fourier component of the stimulus signal (i.e., tagging) as $t$. We assume the following source covariance matrix in the single-stimulus condition (bold elements show that the waveforms of SSVER generators are identical):

$$\mathbf{\Sigma}_{\mathbf{s}}^{\mathbf{1}\text{stim}} = \begin{bmatrix} 1 & \mathbf{1} & 0 & 0 & \\ \mathbf{1} & 1 & 0 & 0 & \\ 0 & 0 & \frac{1}{\kappa} & 0 & \\ 0 & 0 & 0 & \frac{1}{\kappa} & \\ & & & & \cdots \end{bmatrix} \tag{S50}$$

We additionally assume that only SSVER generators are coupled to the stimulus. In the single-stimulus condition, the activity waveforms are assumed to be the same for both generators:

$$\langle s_1 t^* \rangle = \langle s_2 t^* \rangle \equiv c_{st}, \langle |t|^2 \rangle = 1 \tag{S51}$$

$$\langle s_i t^* \rangle = 0, \forall i \notin [1, 2] \tag{S52}$$

Then, the cross-spectrum of the extracted ROI time series and the stimulus can be expressed using the normalized CTF $\tilde{\mathbf{k}}$ as defined in Eq. S15:

$$\langle \hat{r} t^* \rangle = \sum_{i=1}^{N_S} k_i \langle s_i t^* \rangle = k_1 \langle s_1 t^* \rangle + k_2 \langle s_2 t^* \rangle = c_{st} \cdot (k_1 + k_2) \tag{S53}$$

$$\hat{c}_{\hat{r}t} = \frac{\langle \hat{r} t^* \rangle}{\sqrt{\langle |t|^2 \rangle \cdot \langle |\hat{r}|^2 \rangle}} = \frac{c_{st} \cdot (k_1 + k_2)}{\sqrt{\mathbf{k} \boldsymbol{\Sigma}_{\mathbf{s}}^{\mathbf{1}\text{stim}} \mathbf{k}^T}} = c_{st} \cdot \left( \tilde{k}_1 + \tilde{k}_2 \right) \tag{S54}$$

In the two-stimuli condition, we assume a $\pi/2$ delay between the time courses of activity of SSVER generators (and hence zero covariance, as highlighted in bold below):

$$\boldsymbol{\Sigma}_{\mathbf{s}}^{\mathbf{2}\text{stim}} = \begin{bmatrix} 1 & \mathbf{0} & 0 & 0 & \\ \mathbf{0} & 1 & 0 & 0 & \\ 0 & 0 & \frac{1}{\kappa} & 0 & \\ 0 & 0 & 0 & \frac{1}{\kappa} & \\ & & & & \cdots \end{bmatrix} \tag{S55}$$

The predicted ImCoh between ROIs $i$ and $j$ can be calculated using Eq. S22 with $\boldsymbol{\Sigma}_{\mathbf{s}}^{\mathbf{2}\text{stim}}$ in the normalization:

$$\text{Im}(\hat{c}_{ij}^r) = \text{Im}(c_{12}^s) \cdot \left( \tilde{k}_{i1} \tilde{k}_{j2} - \tilde{k}_{i2} \tilde{k}_{j1} \right) \tag{S56}$$

In Fig. S4, we show the predictions of the CTF-based model for an exemplary subject and pipeline (eLORETA followed by averaging), using a $\kappa$ of 100 for illustration purposes. When fitting the models, we performed a grid search over five values of $\kappa$ (10, 20, 50, 100, and 200).

### E. Additional figures and tables

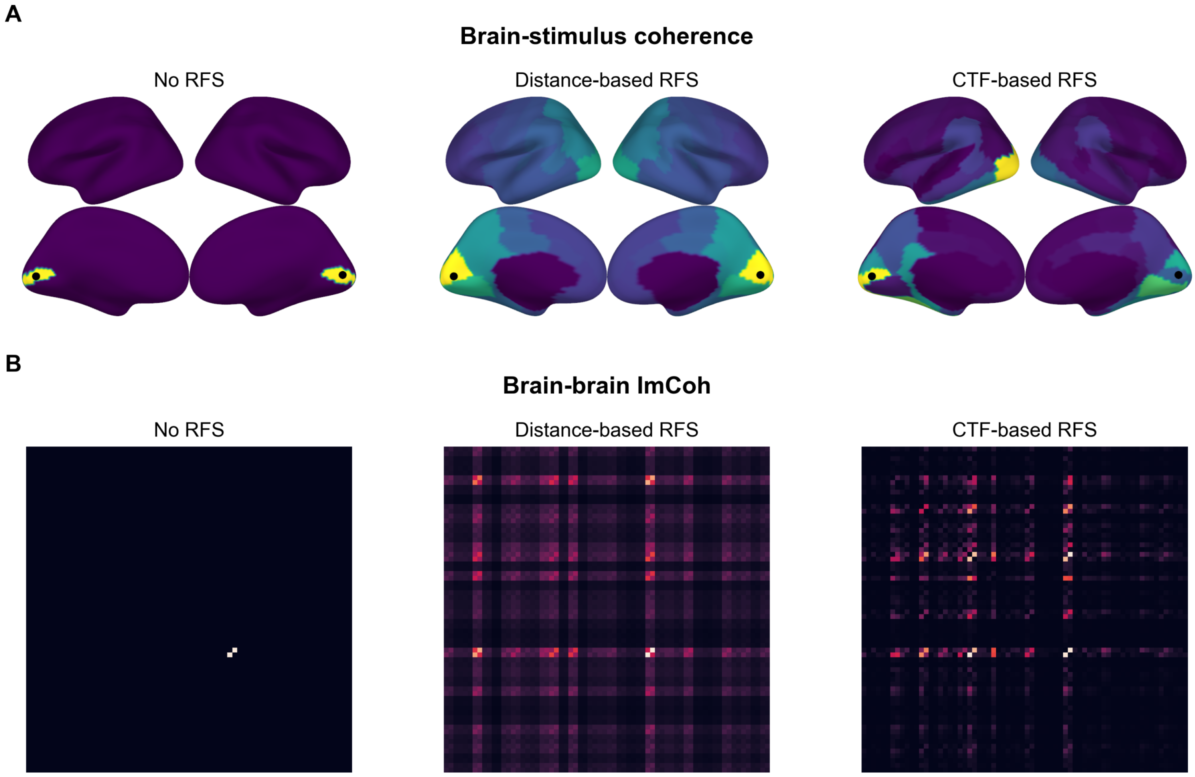


Figure S4: Predictions of the considered models of RFS for exemplary locations of SSVER generators (centroids of the pericalcarine cortex of both hemispheres, defined according to the DK parcellation). For the CTF-based model, the $\kappa$ of 100 was used. (A) Predicted values of brain-stimulus coherence in the single-stimulus condition. (B) Predicted values of brain-brain ImCoh in the two-stimuli condition.

| Comparison | Model | Condition | Correlation | | | |
|---|---|---|---|---|---|---|
| | | | mean | mean-flip | centroid | average |
| raw | CTF | EC | 0.87 | 0.87 | 0.87 | 0.87 |
| | | EO | 0.88 | 0.88 | 0.89 | 0.88 |
| | Distance | EC | 0.62 | 0.61 | 0.69 | 0.64 |
| | | EO | 0.65 | 0.64 | 0.72 | 0.67 |
| delta | CTF | EC | 0.75 | 0.77 | 0.75 | 0.76 |
| | | EO | 0.77 | 0.78 | 0.76 | 0.77 |

Table S3: Correlations between CTF-/distance-based estimates of spurious coherence (SC) and values of SC derived from surrogate data for each extraction pipeline and condition. The right-most column shows the average correlation for the three considered pipelines.

| Comparison | Tagging type | Stimulus phase | Correlation | | |
|---|---|---|---|---|---|
| | | | No RFS | Distance | CTF |
| raw | 1 | fixed | 0.41 | 0.86 | 0.82 |
| | | random | 0.40 | 0.85 | 0.80 |
| | 4 | fixed | 0.42 | 0.81 | 0.79 |
| | | random | 0.45 | 0.81 | 0.79 |
| delta | 1 | fixed | — | — | 0.63 |
| | | random | — | — | 0.60 |
| | 4 | fixed | — | — | 0.72 |
| | | random | — | — | 0.71 |

Table S4: Correlations between the observed values of brain-stimulus coherence and the ones predicted by three considered models of RFS (no RFS, distance-based, and CTF-based). Rows correspond to different experimental conditions of the RIFT dataset (defined by tagging type and the starting phase of the stimulus) and to two types of comparisons performed (raw/delta).

| Comparison | Tagging type | Correlation | | |
|---|---|---|---|---|
| | | No RFS | Distance | CTF |
| raw | 1 | 0.04 | 0.63 | 0.62 |
| | 4 | 0.01 | 0.55 | 0.61 |
| delta | 1 | — | — | 0.36 |
| | 4 | — | — | 0.39 |

Table S5: Correlations between the observed values of brain-brain ImCoh and the ones predicted by three considered models of RFS (no RFS, distance-based, and CTF-based). Rows correspond to different experimental conditions of the RIFT dataset (defined by tagging type) and to two types of comparisons performed (raw/delta).

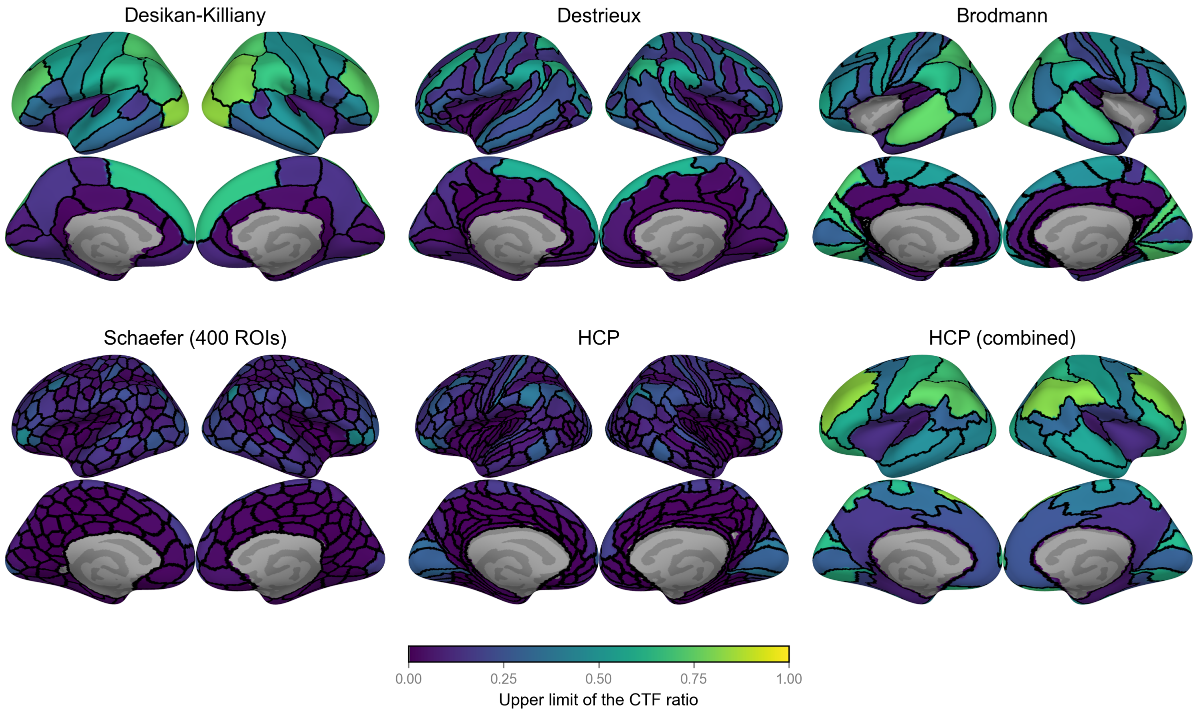


Figure S5: The theoretical upper limits of the CTF ratio for commonly used parcellations. Abbreviations: HCP – Human Connectome Project.

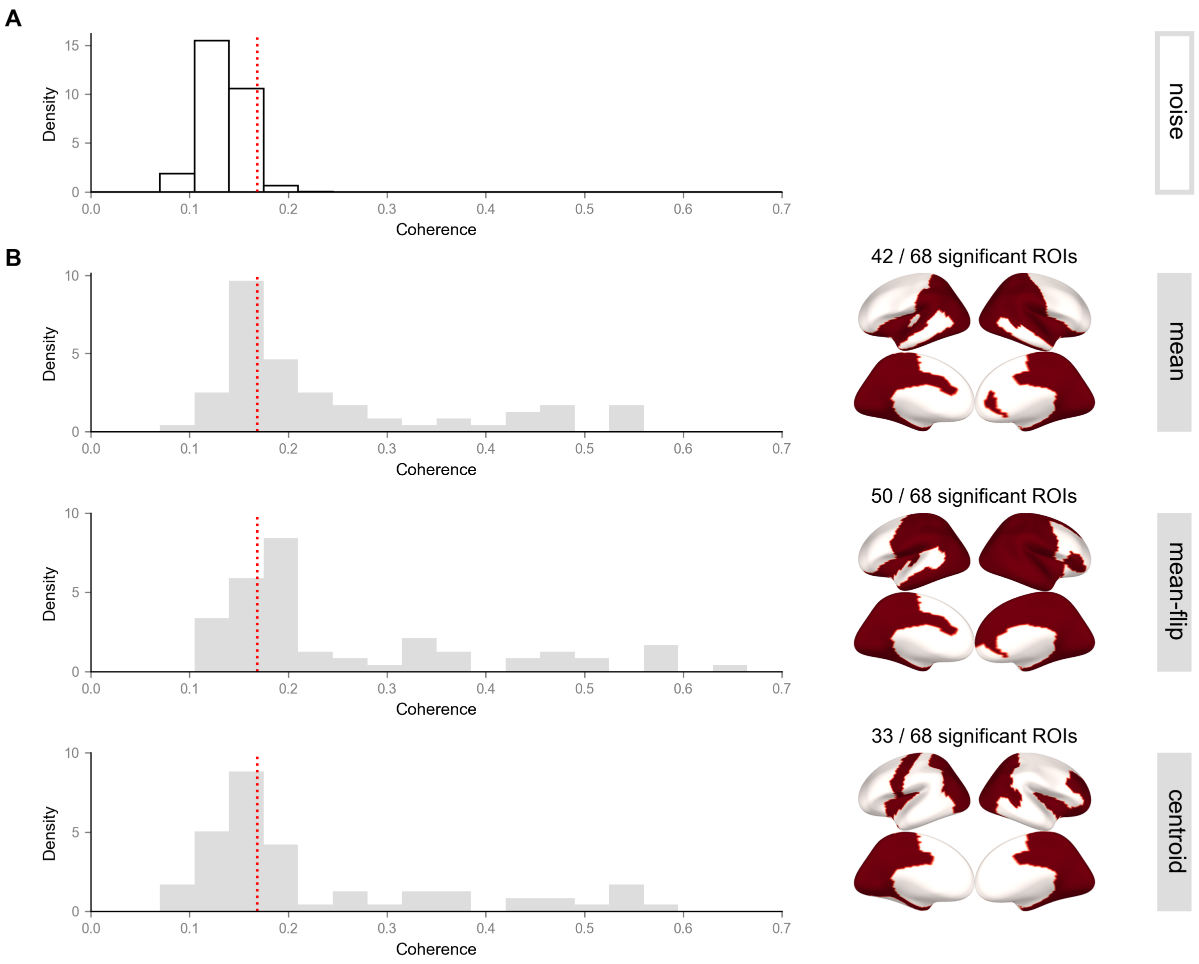


Figure S6: Noise floor for brain-stimulus coherence. (A) Null distribution of coherence between 1000 simulated independent time series. Red dashed line corresponds to the 95th percentile of the distribution, which was used as the threshold for significance. (B) Histograms of brain-stimulus coherence obtained from real data (tagging type 1, fixed starting phase) for each pipeline, and brain masks showing which ROIs exceeded the noise floor.

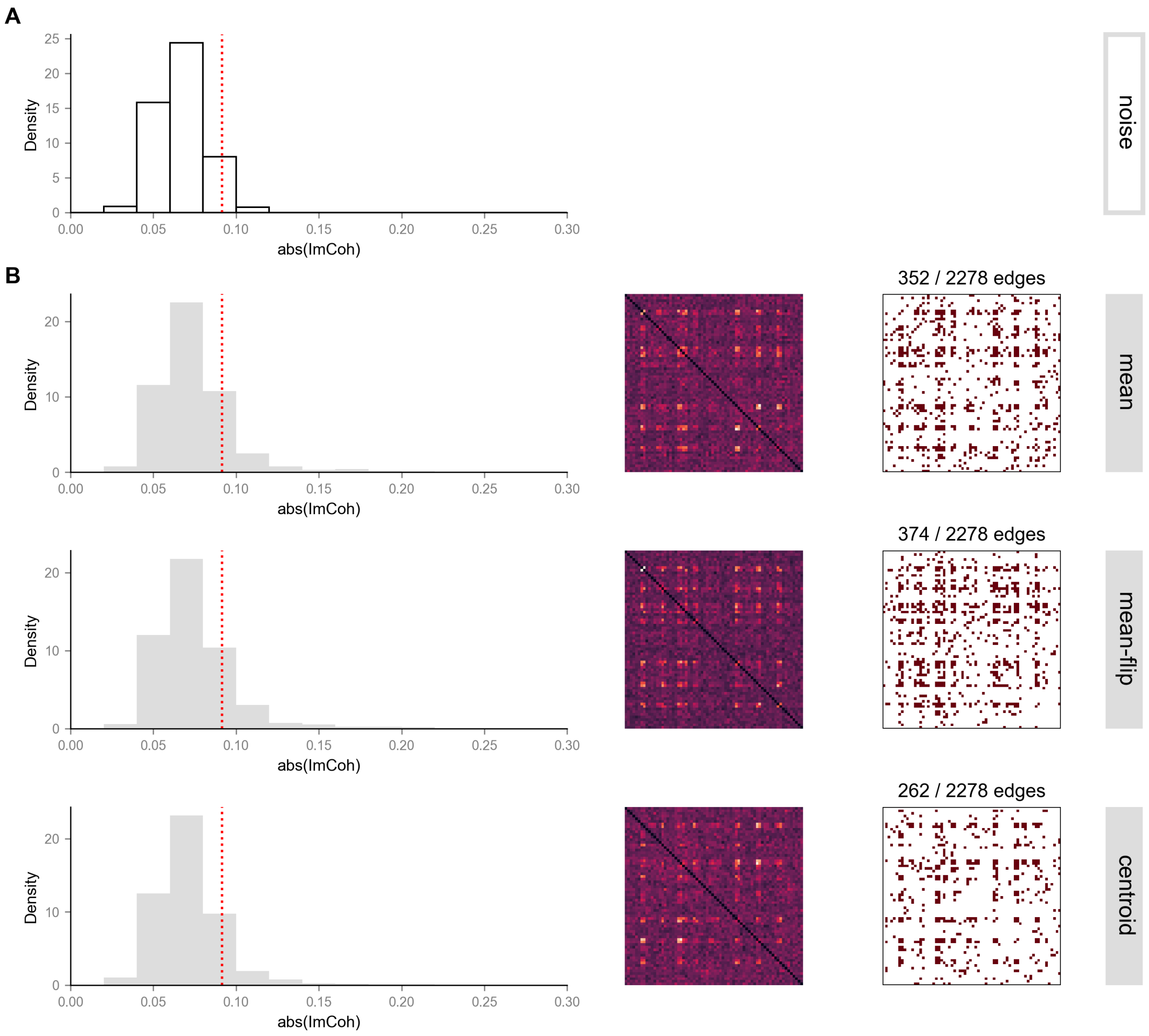


Figure S7: Noise floor for brain-brain ImCoh. (A) Null distribution of the absolute values of ImCoh between 1000 simulated independent time series. Red dashed line corresponds to the 95th percentile of the distribution, which was used as the threshold for significance. (B) Brain-brain ImCoh obtained from real data (tagging type 1, random starting phase) for each pipeline, shown from left to right as a histogram, a connectivity matrix, and a mask indicating brain-brain ImCoh values exceeding the noise floor.

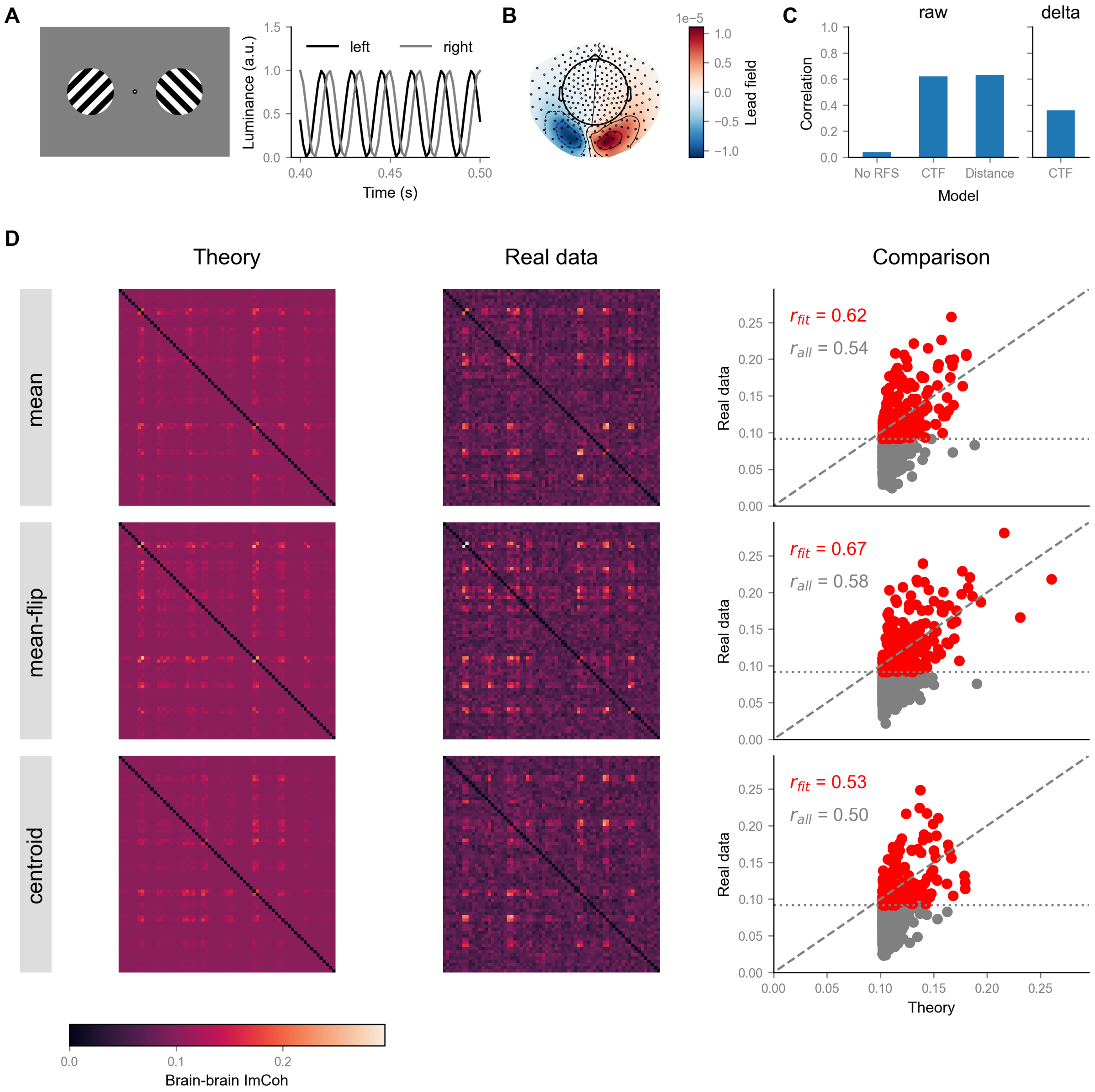


Figure S8: Whole-brain patterns of ImCoh between extracted ROI time series (brain-brain ImCoh) can be explained by two point-like sources located in V1 of both hemispheres using CTF. (A) In the two-stimuli passive viewing condition of the RIFT dataset, there was a $\pi/2$ phase delay between time courses of luminance modulation for the presented stimuli. (B) The lead field of the dipole configuration that yielded the best fit for the CTF-based model. (C) CTF- and distance-based approaches explained comparable variance in the raw values of brain-brain ImCoh, but CTF also partially explained the difference between pipelines for ROI extraction (delta). The correlation values were computed across pooled data from three data-independent extraction pipelines. (D) Theoretical predictions based on CTF and estimates of brain-brain ImCoh from real data are positively correlated for all considered pipelines. For all panels, we used the data from the experimental condition with full luminance modulation (tagging type 1) and random starting phase.

Grosse-Wentrup, M., Liefhold, C., Gramann, K., Buss, M., 2009. Beamforming in noninvasive brain-computer interfaces. IEEE Trans Biomed Eng 56, 1209–19. doi:10.1109/TBME.2008.2009768.

Hashemi, A., Cai, C., Kutyniok, G., Müller, K.R., Nagarajan, S.S., Haufe, S., 2021. Unification of sparse Bayesian learning algorithms for electromagnetic brain imaging with the majorization minimization framework. Neuroimage 239, 118309. doi:10.1016/j.neuroimage.2021.118309.

Haufe, S., Ewald, A., 2019. A Simulation Framework for Benchmarking EEG-Based Brain Connectivity Estimation Methodologies. Brain Topogr 32, 625–642. doi:10.1007/s10548-016-0498-y.

Klein, A., Tourville, J., 2012. 101 labeled brain images and a consistent human cortical labeling protocol. Front Neurosci 6, 171. doi:10.3389/fnins.2012.00171.

Korhonen, O., Palva, S., Palva, J.M., 2014. Sparse weightings for collapsing inverse solutions to cortical parcellations optimize M/EEG source reconstruction accuracy. J Neurosci Methods 226, 147–160. doi:10.1016/j.jneumeth.2014.01.031.

Liljeström, M., Stevenson, C., Kujala, J., Salmelin, R., 2015. Task- and stimulus-related cortical networks in language production: Exploring similarity of MEG- and fMRI-derived functional connectivity. Neuroimage 120, 75–87. doi:10.1016/j.neuroimage.2015.07.017.

Pascual-Marqui, R.D., 2002. Standardized low-resolution brain electromagnetic tomography (sLORETA): technical details. Methods Find Exp Clin Pharmacol 24 Suppl D, 5–12.

Pascual-Marqui, R.D., 2007. Discrete, 3D distributed, linear imaging methods of electric neuronal activity. Part 1: exact, zero error localization. doi:10.48550/ARXIV.0710.3341.

Schaefer, A., Kong, R., Gordon, E.M., Laumann, T.O., Zuo, X.N., Holmes, A.J., Eickhoff, S.B., Yeo, B.T.T., 2018. Local-Global Parcellation of the Human Cerebral Cortex from Intrinsic Functional Connectivity MRI. Cereb Cortex 28, 3095–3114. doi:10.1093/cercor/bhx179.

Scheer, H.J., Sander, T., Trahms, L., 2006. The influence of amplifier, interface and biological noise on signal quality in high-resolution EEG recordings. Physiol Meas 27, 109–17. doi:10.1088/0967-3334/27/2/002.

Timmer, J., König, M., 1995. On generating power law noise. Astronomy and Astrophysics 300, 707.

Tzourio-Mazoyer, N., Landeau, B., Papathanassiou, D., Crivello, F., Etard, O., Delcroix, N., Mazoyer, B., Joliot, M., 2002. Automated anatomical labeling of activations in SPM using a macroscopic anatomical parcellation of the MNI MRI single-subject brain. Neuroimage 15, 273–89. doi:10.1006/nimg.2001.0978.

Van Veen, B., Van Drongelen, W., Yuchtman, M., Suzuki, A., 1997. Localization of brain electrical activity via linearly constrained minimum variance spatial filtering. IEEE Transactions on Biomedical Engineering 44, 867–880. doi:10.1109/10.623056.

Welch, P., 1967. The use of fast Fourier transform for the estimation of power spectra: A method based on time averaging over short, modified periodograms. IEEE Transactions on Audio and Electroacoustics 15, 70–73. doi:10.1109/TAU.1967.1161901.
